# Plastid genome evolution in leafless members of the orchid subfamily Orchidoideae, with a focus on *Degranvillea dermaptera*

**DOI:** 10.1101/2023.11.03.565540

**Authors:** Craig F. Barrett, Matthew C. Pace, Cameron W. Corbett

## Abstract

**Premise:** Leafless, heterotrophic plants are prime examples of organismal modification, the genomic consequences of which have received considerable interest. In particular, plastid genomes (plastomes) are being sequenced at a high rate, allowing continual refinement of conceptual models of reductive evolution in heterotrophs. Yet, numerous sampling gaps exist, hindering the ability to conduct comprehensive phylogenomic analyses in these plants.

**Methods:** We sequenced and analyzed the plastome of *Degranvillea dermaptera*, a rarely collected, leafless orchid species from South America about which little is known, including its phylogenetic affinities.

**Key Results:** We revealed the most reduced plastome sequenced to date among the orchid subfamily Orchidoideae. *Degranvillea* has lost the majority of genes found in leafy autotrophic species, is structurally rearranged, and has similar gene content to the most reduced plastomes among the orchids. We found strong evidence for the placement of *Degranvillea* within the subtribe Spiranthinae using models that explicitly account for heterotachy, or lineage-specific evolutionary rate variation over time. We further found evidence of relaxed selection on several genes and correlations among substitution rates and several other “traits” of the plastome among leafless members of orchid subfamily Orchidoideae.

**Conclusions:** Our findings advance knowledge on the phylogenetic relationships and paths of plastid genome evolution among the orchids, which have experienced more independent transitions to heterotrophy than any other plant family. This study demonstrates the importance of herbarium collections in comparative genomics of poorly known species of conservation concern.

## INTRODUCTION

Leafless, heterotrophic plants provide opportunities to study the genomic consequences of extreme changes in nutritional mode, morphology, physiology, ecology, and evolutionary trajectories (e.g. Bidartondo, 2005; Westwood et al., 2010; Wickett et al., 2014; Těšitel, 2016; Twyford, 2018; Wicke and Naumann, 2018; Yuan et al., 2018; Cai, 2023; Sanchez-Puerta et al., 2023; Timilsena et al., 2023). Plastid genomes have been the focus of intense research over the past few decades, due to their high copy number per cell, relatively small sizes (kilobases for plastomes vs. megabases for nuclear genomes), role in photosynthetic function, and ease with which they can be sequenced via high-throughput technologies (Wicke et al., 2011; Jansen and Ruhlman, 2012; Ruhlman and Jansen, 2014; Twyford and Ness, 2017; Doyle, 2022). The increasing availability of plastid genomes across the plant tree of life has allowed comparative genomic analyses of heterotrophy across multiple taxonomic scales including within species in some cases (Graham et al., 2017; Barrett et al., 2018; Givnish et al., 2018; Wicke and Naumann, 2018; Barrett et al., 2019).

This has allowed a series of related conceptual models of plastome evolution in heterotrophic plants to be developed, with each including sequential refinements as additional plastid genomic resources become available (Wicke et al., 2011; Barrett and Davis, 2012; Barrett et al., 2014; Wicke et al., 2016; Graham et al., 2017; Wicke and Naumann, 2018). The common thread of these conceptual models is that genes become irreversibly pseudogenized and eventually lost in an order more or less associated with their importance in basic plastid function, albeit with many idiosyncrasies among lineages. Genes directly or indirectly involved in photosynthesis tend to be lost first (e.g. NADPH Dehydrogenase, Photosystem, RuBisCO Large subunit, ATP Synthase, Plastid-encoded RNA Polymerase), followed by a series of “housekeeping” genes involved in translation and other basic organellar functions (Maturase K, ribosomal protein genes, transfer RNA genes). Among the last genes to be lost are a set of five “core nonbioenergetic genes” [*accD, clpP, trnE^UUC^, ycf1*, and *ycf2*; *sensu* Graham et al. (2017)], and in a few cases wholescale loss of the plastid genome has been reported [e.g. the endoparasitic angiosperm *Rafflesia* R. Br. (Molina et al., 2014) and the green alga *Polytomella* Aragão (Smith and Lee, 2014)]. Other changes in genomic features frequently occur simultaneously with gene loss, including: changes in GC content; accumulation of repetitive DNA; loss or modification of the Inverted Repeat, genomic deletions, inversions, and rearrangements; and overall increases in substitution rates (Wicke et al., 2016; Barrett et al., 2018; Wicke and Naumann, 2018; Barrett et al. 2019).

Plastid genomes, and genomic datasets more broadly, are increasingly available for phylogenetic placement of heterotrophic lineages, which have long represented some of the most challenging taxa from a systematics perspective due to their reduced morphological, genomic, and sometimes reproductive features (Givnish et al., 2018; Lam et al., 2018; Klimpert et al., 2022). Yet, numerous sampling gaps exist among leafless, heterotrophic lineages throughout the plant tree of life, hindering efforts to produce comprehensive phylogenetic hypotheses and analyses of the process and timing of plastome evolution. While plastomes for some leafless lineages are richly represented, others are poorly sampled or missing altogether, largely due to their diminutive (sometimes cryptic) nature, lack of collections, rarity, and remote habitats (Merckx et al., 2013a, 2013b, Taylor et al., 2013).

Orchids contain more leafless, mycoheterotrophic species than any other plant family. Their requirement of symbiotic fungal germination and subsequent dependence as ‘initial mycoheterotrophs’ upon soil fungi are hypothesized to have facilitated multiple, independent shifts to leaflessness and eventually full mycoheterotrophy with the loss of photosynthetic ability (Leake, 1994; Rasmussen, 1995; Bidartondo, 2005; Freudenstein and Barrett, 2010; Rasmussen et al., 2015; Freudenstein et al., 2017). The leafless/reduced-leaf growth form is predominantly a phenomenon observed in terrestrial orchids, with a few notable exceptions including vining species of the subfamily Vanilloideae Szlach. (e.g. *Erythrorchis* Blume*, Galeola* Lour.) or genera with both terrestrial and epiphytic/lithophitic species (e.g. *Cymbidium* Sw.*, Dipodium* R. Br.; e.g. Motomura et al., 2010; O’Byrne, 2014; Kim et al., 2019). Most leafless orchid species belong to two subfamilies: Epidendroideae Lindl. ex Endl. and Orchidoideae A.A. Eaton, which together constitute >98% of all orchid species diversity (Rasmussen, 1995; Merckx et al., 2013a; Chase et al., 2015; Freudenstein and Chase, 2015; Ogura-Tsujita et al., 2021). The Epidendroideae, which comprise >83% of all species in the family, are predominantly epiphytic, though nearly all leafless or reduced-leaf epidendroids are terrestrial (Freudenstein and Chase, 2015). However, subfamily Orchidoideae (∼15% of all orchid species) is predominantly terrestrial, with centers of diversity in Australia (e.g. *Caladenia* R. Br.*, Corybas* Schltr.*, Pterostylis* R. Br.), Africa (e.g. *Disa* P.J. Bergius), the boreo-temperate Northern hemisphere (e.g. *Goodyera* R. Br., *Platanthera* Rich.), the Pan-tropics (e.g. *Habenaria* Willd.), and the American tropics (e.g. *Cyclopogon* Szlach.*, Microchilus* C. Presl). Leafless or reduced-leaved species in subfamily Orchidoideae are found in several genera within three of the four currently recognized tribes: 1) Orchideae Small (*Brachycorythis* Lindl.*, Platanthera,* and *Silvorchis* J.J. Sm.); 2) Diurideae Endl. ex Butzin (*Arthrochilus* F. Muell.*, Burnettia* Lindl.*, Corybas, Cryptostylis* R. Br.*, Rhizanthella* R.S Rogers, and *Stigmatodactylus* Maxim. ex Makino); and 3) Cranichideae Pfeiff. (*Aspidogyne* Garay*, Chamaegastrodia* Makino & F. Maek.*, Cystorchis* Blume*, Danhatchia* Garay & Christenson*, Degranvillea* Determann, and *Odontochilus* Blume) (Pridgeon et al., 2003; Freudenstein and Barrett, 2010; Merckx et al., 2013a).

To date, complete plastomes have been published for five leafless species within subfamily Orchidoideae: *Corybas cryptanthus* Hatch (tribe Diurideae, subtribe Acianthinae Schltr.; Murray, 2019), *Rhizanthella gardneri* R.S. Rogers (tribe Diurideae, subtribe Rhizanthellinae R.S. Rogers; Delannoy et al., 2011), *Danhatchia australis* (Hatch) Garay & Christenson (tribe Cranichideae, subtribe Goodyerinae Klotzsch; Murray, 2019), and two species of *Chamaegastrodia, C. inverta* (W.W. Sm.) Seidenf. and *C. shikokiana* Makino & F. Maek. (tribe Cranichideae, subtribe Goodyerinae; Tu et al., 2021). Plastomes in these leafless taxa range from 149 kb (*Danhatchia*) to less than 60 kb (*Rhizanthella*), and display a range of gene loss. Yet, these sequences represent only two tribes and three subtribes among the orchidoids (of the eight orchidoid subtribes containing leafless taxa), leaving several important sampling gaps in this subfamily of predominantly terrestrial orchids.

One example is *Degranvillea dermaptera* Determann (Fig. 1). This species, to our knowledge, has only been collected a handful of times. The Global Biodiversity Information Facility (GBIF) lists 18 preserved specimens, held in just a few herbaria, and 12 human observations in the form of iNaturalist records (GBIF.org, last accessed 11 October 2023). Little is known about this species, which is endemic to French Guiana in South America, and was described in the mid-1980s (Determann, 1985; Merckx et al., 2013a). While this species has been included in several checklists or mentioned in floristic summaries of the Guyana Region, no formal conservation assessment has been conducted (Dumont et al., 1996; Bikaeff, 2002; Funk et al., 2007; Véron et al., 2021). The most recent, comprehensive orchid taxonomic treatment of Chase et al. (2015) placed the genus within the subtribe Spiranthinae (tribe Cranichideae), which predominantly share a terrestrial habit. The Spiranthinae are well studied in terms of morphological variation and phylogenetic relationships based on molecular markers (e.g. Garay, 1982; Burns-Balogh and Robinson, 1983; Salazar et al., 2003; Gorniak et al., 2006; Borba et al., 2014; Salazar et al., 2018), representing “…the most species-rich clade of terrestrial orchids in the Neotropics, where most of the c. 40 genera and 520 species are found… (Salazar et al., 2018)”. The phylogenetic placement of *Degranvillea*, as with several other features including fungal host associations, plastid genome structure, and conservation status, remain a mystery. Salazar et al. (2018) further state: “…*Degranvillea* shows some floral features in common with the *Pelexia* [Poit. ex Lindl.] clade, including the narrowly conical sepaline spur, subulate nectar glands of the labellum and truncate rostellum with terminal viscidium (as in some species of *Sauroglossum* [Lindl.] and *Pelexia* s.s.), but its highly modified vegetative organs do not offer any obvious clue about its phylogenetic affinities.”

**Fig. 1.**
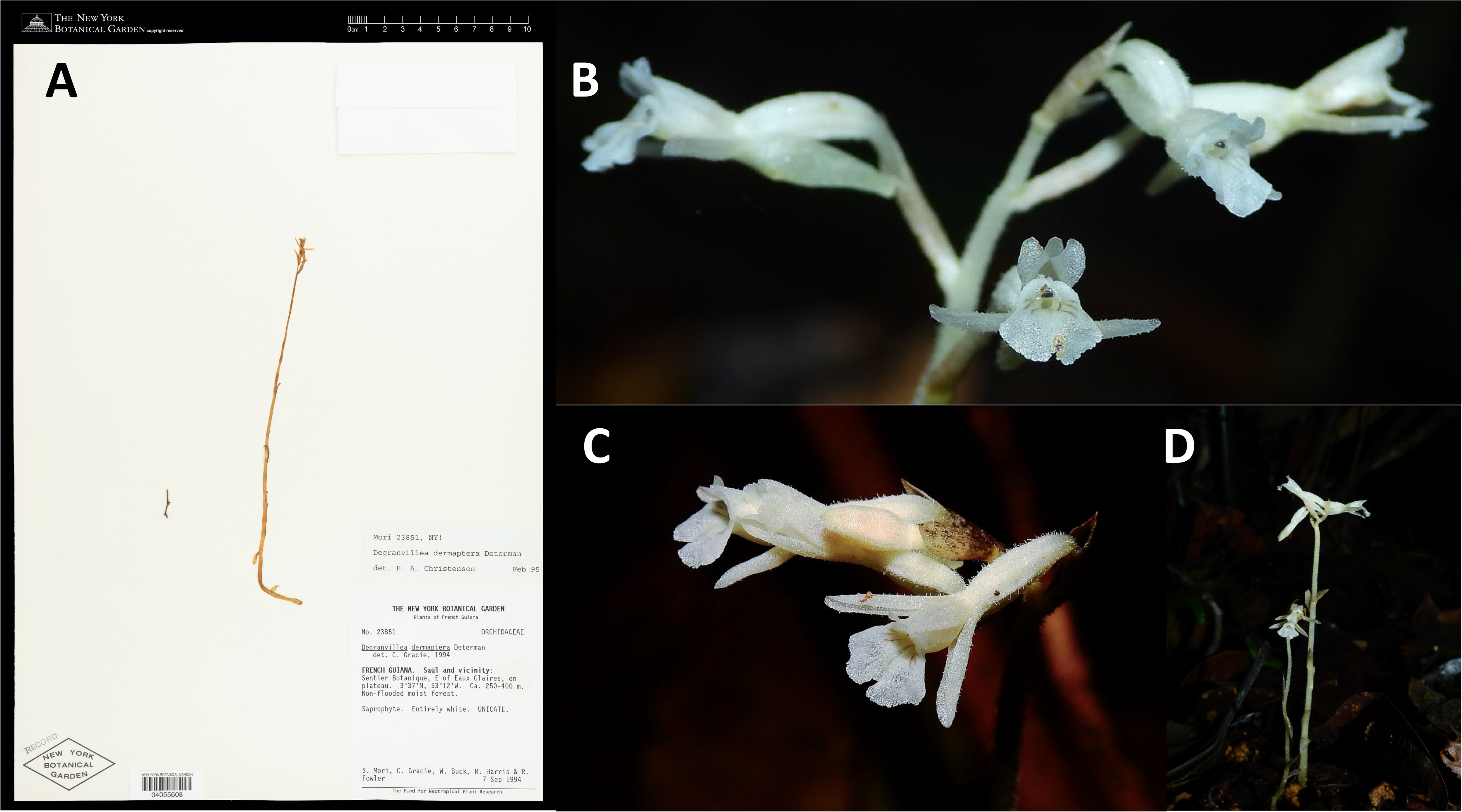
**A**. The sampled specimen of *Degranvillea dermaptera* (Mori number 23851). **B**. and **C**. Close-up of flowers and inflorescence. **D**. Habit. Photos: Image courtesy of the C. V. Starr Virtual Herbarium (http://sweetgum.nybg.org/science/vh/) and Sébastien Sant (Guiana Amazonian Park, https://www.inaturalist.org), with permission.

We sequenced the plastome of *Degranvillea dermaptera*, characterized its structure, compared the degree of genomic reduction to other mycoheterotrophic orchids, and estimated its phylogenetic relationships within subfamily Orchidoideae based on complete plastid genomes and nuclear ITS sequences. *Degranvillea* has the most reduced plastome sequenced to date among subfamily Orchidoideae (though not among orchids more broadly), lacking all photosynthesis-related genes and encoding only “housekeeping” genes involved in translation or a few other nonbioenergetic processes. We applied phylogenetic methods that specifically model heterotachy (i.e. lineage-specific heterogeneous substitution rates over time) to the plastome data and found strong support for the phylogenetic placement of *Degranvillea* among the orchidoid subtribe Spiranthinae Lindl. ex Meisn., within which it is currently circumscribed based on morphology. We analyzed structural variation of the plastome, demonstrating extensive gene losses, relaxed selection pressure for multiple genes among leafless orchidoids, and correlations among substitution rates and other plastid genomic features in a phylo-comparative framework. This study advances knowledge on the phylogenetic affinities and genome evolutionary trajectories in leafless heterotrophs, which serve more broadly as ‘evolutionary experiments’ that are informative on the limits of organellar genomic modification.

## METHODS

### Sampling, DNA extraction, library building, and sequencing

Plant material (0.5 cm^2^) was sampled from the packet of NYBG Steere Herbarium specimen *Mori 23851* (NY). The specimen was collected in French Guiana, with the locality description (Fig. 1A): “Saül and vicinity: Sentier Botanique, E of Eaux Claires, on plateau. 3 37’N, 53 12’W. Ca. 250-400 m. Non-flooded moist forest. Saprophyte. Entirely white. S. Mori, C. Gracie, W. Buck, R. Harris, & R. Fowler. 7 Sep 1994.” Total genomic DNAs were isolated using a 1/10 volume CTAB protocol (Doyle and Doyle, 1987), which was further modified by adding an extra 24:1 chloroform:isoamyl alcohol separation. The DNA pellet was eluted in 120 ul of Tris-EDTA (pH 8.0), and run on a 1% agarose gel to assess the level of DNA degradation. Further, a NanoDrop spectrophotometer (Thermo Fisher, Waltham, Massachusetts, USA) was used to check DNA quality, and a Qubit fluorometer (broad range assay, Thermo Fisher) was used to quantify double stranded DNA. The double stranded gDNA concentration was 3.9 ng/ul. A total of ten ng of DNA was used for Illumina library preparation with the SparQ DNA Frag and Library Kit (Quantabio, Beverly, Massachusetts, USA), in a 20 ul volume (2/5 the standard protocol), and modified in that a 1 min fragmentation time was used instead of 14 min to prevent overshearing. The final library was quantified via Qubit high sensitivity assay and visualized on a 1% agarose gel. Smaller library fragments (< 200 bp) were removed with a 0.8× bead:sample ratio cleanup with Quantabio SparQ PureMag beads. The library was pooled with 63 others at equimolar ratios and library fragment size and concentration was checked on an Agilent Bioanalyzer (Agilent Technologies, Santa Clara, California, USA). The libraries were sequenced on an Illumina NextSeq2000 using v3 chemistry (Illumina, San Diego, California, USA), producing approximately 10^9^ 2×100 bp reads.

### Read processing, plastome assembly, and annotation

Reads were processed with FASTP (Chen et al., 2018), using adapter overhang trimming, a sliding window of 16 bp to remove poly-G and poly-X tails that are common with libraries derived from herbarium specimens containing short fragments, and keeping only reads with a minimum length of 75 bp (-w 16 -m 75 - trim_polyg, -trim_polyx -l 75 -cut_right). Resulting trimmed and filtered FASTQ files were assembled *de novo* with GetOrganelle v. 1.7.7.0 (Jin et al., 2020) and NOVOPlasty v. 4.3.1 (Dierckxsens et al., 2017). *Spiranthes sinensis* (Pers.) Ames (GenBank accession OM314917) was used as a reference/seed sequence for both. *Spiranthes* is in subtribe Spiranthinae, the same subtribe where *Degranvillea* has been hypothesized to be placed (Salazar et al., 2018). The GetOrganelle assembly was conducted specifying a SPADES (Bankevich et al., 2012) k-mer range of 25-85, and using all available reads (-t 28 -R 100 -k 21,31,45,65,85 -F embplant_pt -- max-reads inf --reduce-reads-for-coverage inf). NOVOPlasty was run in a similar fashion, using the same reference/seed, with insert size set to ‘auto’ and a kmer length of 33. Trimmed reads were mapped back to the resulting contigs from Getorganelle and NOVOPlasty in Geneious v.10 under medium/high sensitivity, and allowing ‘find structural variants, short insertions, and deletions of any size’ to check the quality and accuracy of the assemblies. The Inverted Repeat (IR) was detected by searching for exact repeats > 1000 bp in Geneious, and verified with mapped, paired end reads across the IR boundaries. The genome was then manually rearranged in the configuration of Large Single Copy (LSC) > Inverted Repeat B (IR_B_) > Small Single Copy (SSC) > Inverted Repeat A (IR_A_). The reoriented plastomes from both assemblers were then pairwise-aligned with MAFFT v7.271 (Katoh and Standley, 2013) with the ‘-auto’ option. The final plastome was annotated with GeSeq (Tillich et al., 2017), keeping the best annotation from Chlöe (Zhong, 2020), HMMER (http://eddylab.org/software/hmmer/hmmer.org), or BLAT (Kent, 2002), with the latter specifying a 25% similarity threshold to account for highly divergent genes. The annotated GenBank file was then further edited in Geneious, and each open reading frame (ORF) was verified with the Geneious ORF-finder. MapDamage v.2.0 (Jónsson et al., 2013) was use to assess the degree of C/T and G/A DNA damage patterns across both forward and reverse reads. Cleaned reads were mapped to the final reference genome with BBMAP v. 38.51 (http://sourceforge.net/projects/bbmap/) and the resulting .bam file was used as input for MapDamage.

### Genome structure

Genomic rearrangements were analyzed by aligning the *D. dermaptera* genome to that of *Spiranthes sinensis* as a reference with the Geneious plugin for MAUVE (Darling et al., 2010), using MAFFT for the final gapped alignment. Three distance metrics were calculated in MAUVE—double-cut-and-join, single-cut-and-join, and breakpoint—to determine the number of locally collinear blocks between them. Expansions and contractions of the IR were visualized among all leafless taxa and *Spiranthes sinensis* with the IRPlus web application (Menéndez et al., 2023): https://irscope.shinyapps.io/IRplus/. The breakpoints of gene inversions were investigated more closely by extracting the region from the MAUVE alignment in Geneious and searching for palindromic repeats with the PALINDROME web interface to EMBOSS (Rice et al., 2000; https://www.bioinformatics.nl/cgi-bin/emboss/palindrome), specifying perfect repeats, minimum length of 10 bp, and maximum length of 100 bp. Positive motif hits were then added as annotations in Geneious, and the palindrome and intervening sequences were used as a queries in MFOLD (Zuker, 2003) with default parameters via the UNAfold Web Server (http://www.unafold.org/mfold/applications/dna-folding-form.php). A comparison of functional gene content was conducted among representatives of all publicly available leafless orchid genera and a representative leafy orchid from each subfamily. Gene annotations (CDS, rRNA) were extracted from Geneious and converted to a presence/absence matrix in Microsoft Excel (Microsoft Corporation, 2018) as a .csv file. This was imported into R (R Core Team, 2023) and converted into a binary heatmap with pheatmap v.1.0.12 (Kolde, 2019).

### Phylogenomic analyses

Complete plastomes were downloaded for available members of subfamily Orchidoideae, as well as representative outgroups from the four other orchid subfamilies. Also included were the plastid gene alignments from Serna-Sánchez et al. (2021). GenBank flat files were imported into PhyloSuite v.1.2.3 (Zhang et al., 2020), and all CDS and rDNAs were extracted. Data for each locus were combined in Geneious and aligned with MAFFT (-auto). All alignments were checked for anomalous sequences; in the case of *ycf1*, several truncated sequences were identified (partial sequences from the IR), removed, replaced with full length CDS, and realigned. Alignments were concatenated into a single matrix in Geneious, and phylogenetic analysis was conducted with IQTree2 (Minh et al., 2020). Because the dataset contained fully mycoheterotrophic taxa, some of which are known to have drastically accelerated substitution rates, we implemented the GHOST heterotachy model (Crotty et al., 2020). GHOST is an edge-unlinked mixture model, allowing separate rate classes and base frequencies among site classes, thus modeling heterotachy, or rate variation within and among lineages over time. We first implemented ModelFinder (Kalyaanamoorthy et al., 2017), which selected ‘GTR’ among the major model classes. We tested among different numbers of rate classes (“K” = 2, 4, 6, and 8) using the GHOST heterotachy model (-m GTR+FO*H{2,4,6,8}), and compared each model using the Bayesian Information Criterion (BIC) and BIC weights (Schwarz, 1978), with calculations of the latter carried out in R.

A second analysis was conducted using the Internal Transcribed Spacer (ITS) region from the nuclear genome. ITS sequences from Salazar et al. (2018), which featured nearly complete generic level sampling of subtribe Spiranthinae including several outgroup taxa within subfamily Orchidoideae, were downloaded from NCBI GenBank, as well as *Platanthera chlorantha* (Custer) Rchb. f. (GenBank Accession MF944373) as an additional outgroup. Cleaned reads from *Degranvillea* were mapped iteratively to the ITS region of *Cyclopogon congestus* (Vell.) Hoehne (Accession MG360476) in Geneious, under low-medium sensitivity for five iterations. The consensus sequence was then aligned with all other taxa with MAFFT (-auto). The best fit model for the ITS alignment was identified with ModelFinder in IQTree2. We did not apply the GHOST model to ITS data, because the number of parameters quickly exceeds the number of sites given the taxon sample (>300 accessions) for values of K >2, leading to issues with parameter identifiability (Burnham and Anderson, 2002).

Divergence time analyses were conducted via least squares dating (LSD; To et al., 2016), implemented in IQtree2. LSD uses a Gaussian model, closely related to the Fitch-Langley molecular clock model, that has been shown to be robust to uncorrelated violations of the molecular clock, producing similar estimates to current Bayesian and ML methods (e.g. BEAST; Drummond et al., 2007) at a fraction of their computation time, and without the need to designate extensive priors on parameters (Kumar and Hedges, 2016; To et al., 2016). The goal of this analysis was to place a relative time scale on plastome degradation for *Degranvillea* and other leafless taxa, and not to conduct a comprehensive divergence time analysis for Orchidaceae Juss., which has been addressed elsewhere (Givnish et al., 2016; Serna-Sánchez et al., 2021; Zhang et al., 2023; Pérez-Escobar et al., 2023). We used an internal calibration of 15-20 million years with the fossil species *Meliorchis cariba* Ramírez for the crown age of subtribe Goodyerinae (Ramírez et al., 2007), while specifying a root-age range for the orchids at 120-70 million years based on estimates from several previous studies (Givnish et al., 2016; Serna-Sánchez et al., 2021; Zhang et al., 2023; Pérez-Escobar et al., 2023). Because LSD requires a hard calibration when using a single fossil calibration point, we used two calibration times in separate analyses—20 mya and 15 mya—which correspond to the maximum and minimum estimates for *Meliorchis*/Goodyerinae (Ramírez et al., 2007). The ML tree resulting from the GHOST heterotachy analysis was used (GTR+FO*H6), with *Apostasia odorata* Blume as the outgroup, and the same model was used in the LSD analysis (parameters in IQTree2: -o “Apostasia_odorata” -T 28 --date-root -120:-70 --date-options “-u 1 -z 0” --date-ci 1000). An analysis was also conducted for the ITS tree and dataset, as above, but specifying the best-fit model identified by ModelFinder, and calibrating the node corresponding to six members of Goodyerinae (from genera *Dossinia* C. Morren*, Goodyera, Ludisia* a. Rich.*, Pachyplectron* Schltr., and *Platylepis* A. Rich.) with both 20- and 15-my dates. Confidence limits on the age estimates for ancestral nodes were generated via bootstrap resampling, and were visualized with FigTree v.1.44 (Rambaut, 2010). The current approach was taken based on robustness of results in simulations and empirical studies comparable to those of other ML and Bayesian approaches (e.g. Jones and Poon, 2016; To et al., 2016; Miura et al., 2020), and because we were able to use the entire plastid sequence dataset, as model-based (especially Bayesian) approaches can be intractable for phylogenomic datasets, often requiring downsizing to a few genes to reach convergence (e.g. Barrett et al., 2018; Givnish et al., 2018).

### Genome evolutionary analyses

RELAX (Wertheim et al., 2015) in HyPhy v.2.2.4 (Kosakovsky Pond et al., 2005) was used to test for significantly relaxed (ω, or *dN*/*dS* ∼ 1) or intensified selection pressure (*dN*/*dS* > or < 1) for all CDS with intact reading frames in *Degranvillea dermaptera* (Table 1). First, we removed all stop codons in the FASTA alignments and adjusted them to be in reading frame with the codon-aware aligner MACSE v.2 (Ranwez et al., 2018; command: “-prog exportAlignment -align <alignment.fasta> -codonForFinalStop ---”). RELAX requires a codon-based multiple sequence alignment and a phylogenetic tree with the same taxa. We used the tree from the GHOST heterotachy analysis above as our reference tree. Because some genes were missing for taxa other than *Degranvillea*, we read both the tree and the FASTA alignments into R using the *read.tree* and *read.dna* functions in APE v.5.7-1 (Paradis and Schliep, 2019), and used the *name.check* function of GEIGER v.2.0 (Pennell et al., 2014) to verify which taxa in the tree might be missing for each locus. The *drop.tip* function in APE was then used to prune taxa such that the tree could be articulated with each corresponding FASTA alignment. Two rounds of RELAX analyses were conducted: one in which representatives of all five leafless genera were designated as test branches (*Chamaegastrodia inverta, Corybas cryptanthus, Danhatchia australis, Degranvillea dermaptera,* and *Rhizanthella gardneri*), and another in which only *Degranvillea* and *Rhizanthella* were specified, as these two taxa showed the most evidence of plastome modification (see below). RELAX output (.json) was visualized with HyPhy Vision (http://vision.hyphy.org). Commands for data processing and HyPhy analyses can be found at https://github.com/barrettlab/HyPhy-molecular-evolution---RELAX/wiki.

**Table 1.**
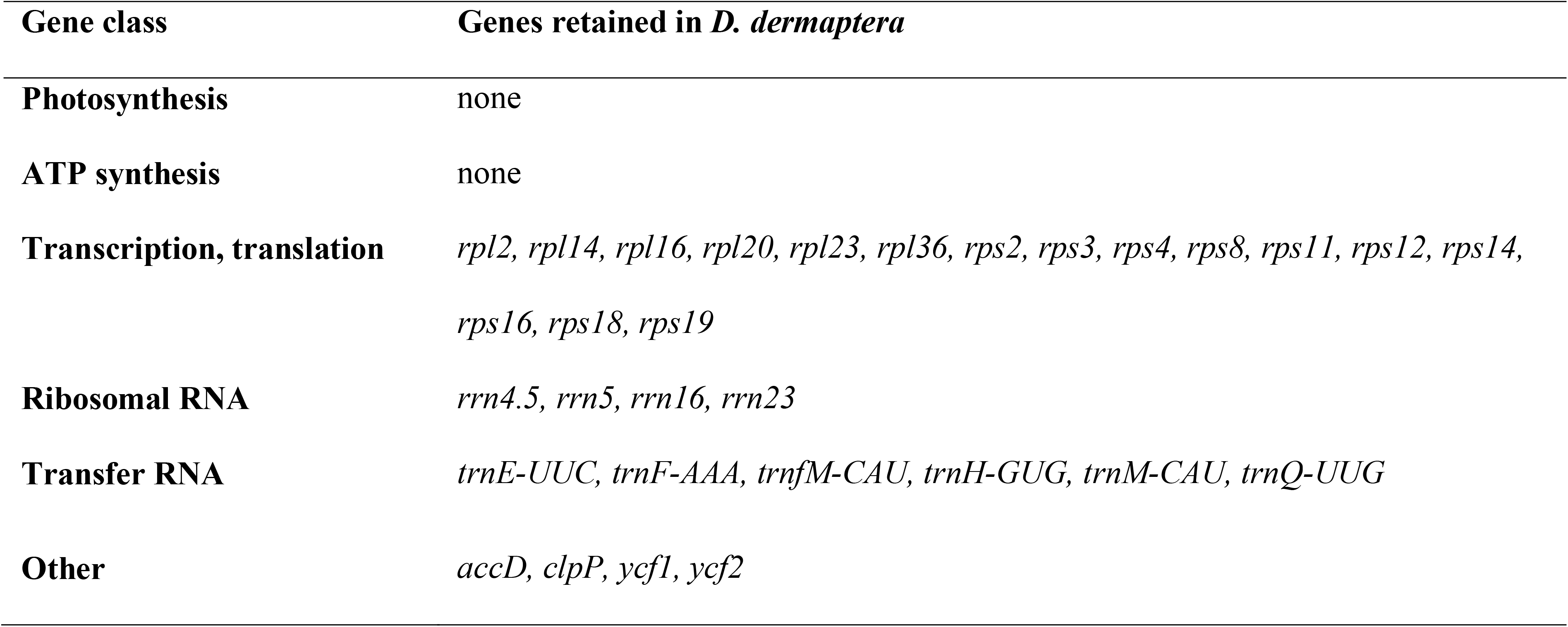
Genes physically retained as putatively functional in *Degranvillea dermaptera*.

A global, phylo-comparative analysis among substitution rates and “traits” of the plastid genome was conducted with CoEvol v.1.6 (Lartillot and Poujol, 2016), using the five representative leafless taxa above and one leafy taxon (*Spiranthes sinensis*). Specifically, correlations were tested among: *dN, dS,* GC content, plastome length, IR length, the number of functional CDS, the number of functional transfer RNAs, mean intron length for intron-containing CDS shared among all taxa, root-to-tip patristic distances, the number of perfect repeats >20 bp, double-cut-and-join distance, and density of microsatellite repeats across the genome. CoEvol conducts Bayesian inference via Markov Chain Monte Carlo methods using a probabilistic, multivariate, Brownian process model on the sequence alignment (here, the codon-based, concatenated alignment of all CDS present in *D. dermaptera* as above), a tree, and a set of “traits.” The model further conducts correlations between all pairs of substitution classes (*dN, dS*) and traits while simultaneously accounting for all other trait correlations. GC content, plastome length, number of CDS, and number of tRNAs were calculated for each of the included plastomes (minus one IR copy) in Geneious. IR length was also calculated in Geneious, as was the number of repeats >20 bp, using the native “find repeats” feature. Root-to-tip patristic distances were calculated with the R packages APE and ADEPHYLO v.1.1-16 (Jombart and Dray, 2008) based on the tree resulting from the GHOST GTR+FO*H6 model (as above), which was pruned with the *drop.tip* function of APE in R. Microsatellite repeat density was calculated as the number of microsatellite repeat regions detected divided by total plastome length (with one IR copy removed). Microsatellite repeats were detected with MISA-web (Beier et al., 2017), specifying the following parameters (SSR motif length/minimum number of repetitions): 1/10, 2/6, 3/5, 4/3, 5/3, and 6/3. Three independent CoEvol chains from random seeds were run for 10^4^ generations with a burn-in of 10^3^ generations, sampling every 10 generations. The ‘tracecomp’ program, part of the CoEvol distribution, was used to assess stationarity and convergence of the chains, requiring effective sample sizes of >300 and ‘maxdiff’ values < 0.1 for each parameter. Final correlation matrices were calculated with the ‘readcoevol’ script, also provided with CoEvol.

## RESULTS

### Genome structure

A total of 43,543,675 paired end reads were generated for *D. dermaptera*, with 36,484,708 remaining after trimming with FASTP. Mean coverage depth of the plastome was 101.6×, with a mean insert size of 181.4 bp. Both GetOrganelle and NOVOPlasty produced identical, circularized chromosomes of 46,846 bp (Fig. 2). The genome contains 20 protein coding DNA sequences (CDS), four ribosomal RNA genes (rDNA), and six transfer RNA genes (tRNA) (Table 1; Fig. 2). Five genes retain their introns: *3’-rps12*, *rpl2*, *rpl16*, *rps16*, and *clpP*, the latter with two introns present. Two of these, *rpl2* and the 3′L*rps12* (exons 2 and 3), are Group IIA introns.

**Fig. 2.**
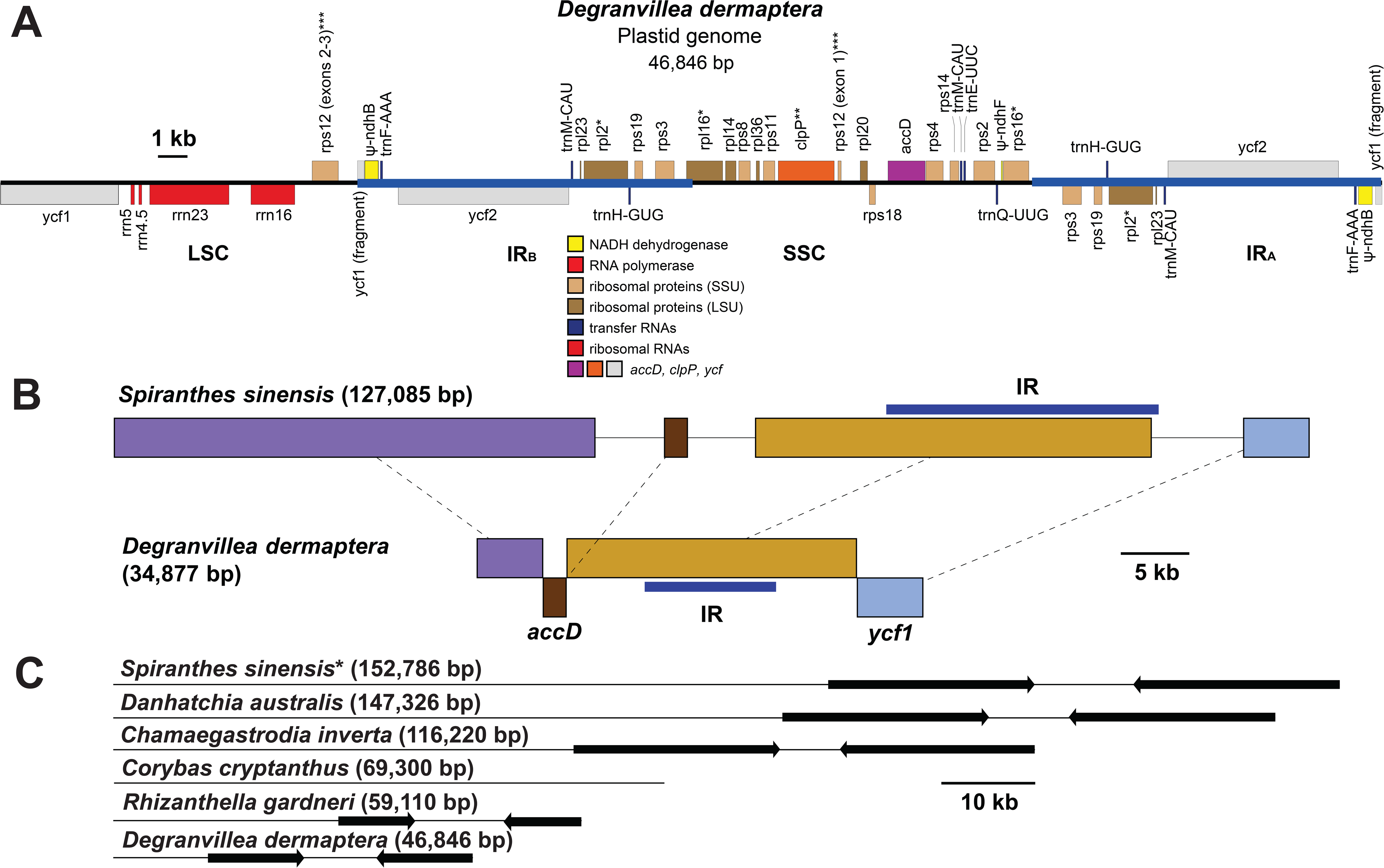
**A**. The plastid genome of *Degranvillea dermaptera*, in linear format. Scale bar = 1 kilobase (kb). ‘*’ indicates that a particular gene contains one intron; ‘**’ indicates two introns; ‘***’ refers to *rps12*, which contains one intron between cis-spliced exons 2 and 3, and trans-spliced exon 1; ‘_J’ indicates a pseudogene. Blue horizontal bars indicate the Inverted Repeat (IR). **B**. Locally colinear blocks between the plastomes of *Spiranthes* and *Degranvillea* with one IR copy removed. Scale bar = 5 kb. Blue horizontal bar indicates the IR. Inverted segments in *Degranvillea* indicate major inversions of the plastomes. Colors correspond to syntenic locally colinear blocks. **C**. Scaled length comparison among the leafy *Spiranthes sinensis* (*) and five leafless taxa (below). Scale bar = 10 kb. Black arrows indicate the IR.

Notable genes missing in *D. dermaptera* are all Photosystem I and II genes (*psa*, *psb*), all ATP Synthase genes (*atp*), the RuBisCo Large Subunit gene (*rbcL*), the Maturase K gene (*matK*), and all Cytochrome b/f Complex genes (*pet*). All NADPH Dehydrogenase genes are also missing, except for small fragments of ψ-*ndhB* and ψ-*ndhF*, which are both truncated pseudogenes. Of the 20 CDS remaining in the plastome of *D. dermaptera*, most are ribosomal proteins, with the addition of *accD*, *clpP, ycf1*, and *ycf2*. Three of the latter four CDS are significantly shorter than their homologs in *S. sinensis*: *accD* (1263 bp in *D. dermaptera* vs. 1482 bp in *S. sinensis*); *ycf1* (4204 bp vs. 5469 bp), and *ycf2* (5787 bp vs. 6651 bp), while *clpP* is 588 bp in both species. The six remaining tRNA genes are *trnE^UUC^*, *trnF^AAA^, trnfM^CAU^, trnH^GUG^, trnM^CAU^*, and *trnQ^UUG^*. A comparison of functional plastid gene content among published orchid plastomes revealed that *Degranvillea* is in the ‘advanced’ stages of degradation, with similar gene content to the most reduced orchid taxa, including *Epipogium*, *Gastrodia*, *Rhizanthella*, and *Pogoniopsis* (Fig. 3). Analysis of DNA damage patterns with MapDamage revealed negligible effects of C/T or G/A base misincorporation (Appendix S1). C/T and G/A misincorporation rates were highest at the 5’ ands of both the forward and reverse reads mapped to the plastome, respectively, which rapidly decreased along the lengths of each read, and never exceeded substitution rates of > 0.012.

**Fig 3.**
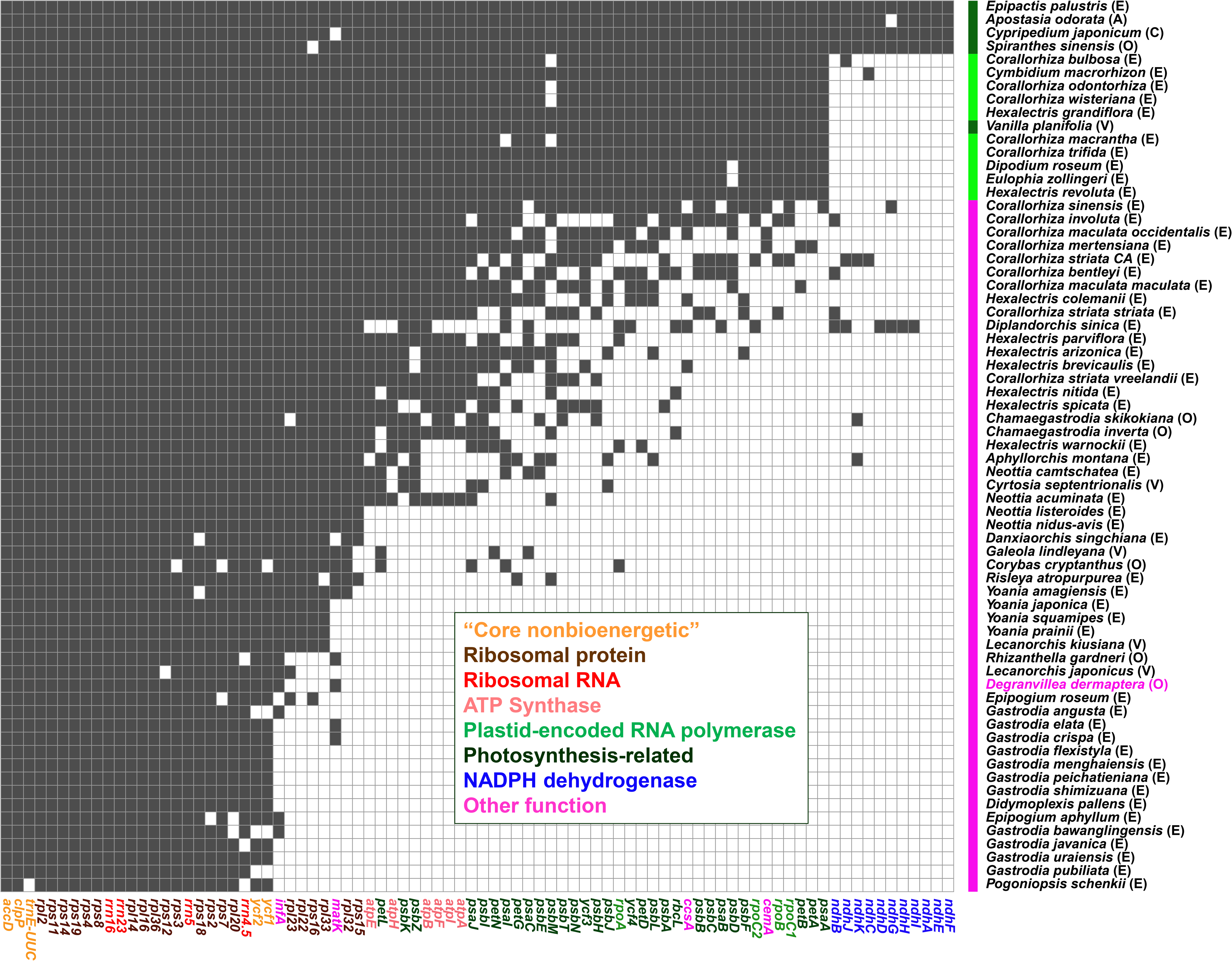
Comparison of functional gene content among leafy (right bar, dark green), leafless, putatively partial mycoheterotrophs (right bar, light green), and leafless full mycoheterotrophs (right bar, magenta). Black squares indicate retention of a functional gene and white squares indicate absence or a pseudogene. Letters adjacent to species names indicate subfamilies Epidendroideae (E), Orchidoideae (O), Cypripedoideae (C), Vanilloideae (V), and Apostasioideae (A). Genes are colored by functional class along the horizontal axis, with “Core nonbioenergetic” genes *accD, clpP, trnE^UUC^, ycf1*, and *ycf2* colored in orange (*sensu* Graham et al., 2017).

The IR region is 11,969 bp in length, and the LSC and SSC are 12,007 and 10,901 bp, respectively (Fig. 2A). The SSC region contains genes that are typically found in the LSC of other land plants (e.g. *accD*, *clpP*), thus likely representing part of the LSC ancestrally. The *rpl16* gene spans the IR_A_-LSC boundary, the *rps16* intron spans the IR_B_-SSC boundary, and the second exon of *rps16* terminates within the SSC just before the SSC-IR_A_ boundary (Appendix S2). Genome structural analysis in MAUVE comparing *D. dermaptera* to *S*. *sinensis* (GenBank accession OM314917; Fig. 2B) revealed two major genomic rearrangements, with a double-cut-and-join distance (DCJ) of 2, a single cut and join distance of 6, and a breakpoint distance of 3. MAUVE recovered four locally colinear blocks (LCB, aligned lengths given): 1) 5 kb, consisting of part of the *ycf1* gene; 2) 43 kb, consisting of a syntenic region spanning *rpl20* to *rrn4.5*; 3) 1 kb, consisting of a part of *accD*; and 4) 41 kb, consisting of a syntenic region from *rps16* to *rps4*. A closer investigation of the *accD* inversion revealed a relatively high density of palindromic repeats near the gene boundaries that are not present in *S*. *sinensis* (Appendix S3). Three of these coincide with putative breakpoints of the *accD* inversion. The first revealed a cruciform structure near the 5’ end of *accD* (palindrome 287, 123 bp in total, motif length = 10 bp, deltaG = -8.87). This same motif was also found in inverted orientation just beyond the 3’ terminus of *accD*. The second palindrome was a hairpin-loop structure in the same region (p350, 90 bp in total, motif length = 10 bp, deltaG = -7.87). The third formed a hairpin just at the 3’ end of *accD* (palindrome 1639, 52 bp in total, motif length = 19 bp, deltaG = -16.95).

Comparison of the five leafless species included within subfamily Orchidoideae with the leafy *Spiranthes sinensis* (152,786 bp) revealed a range in the degree of genome size reduction (Fig 2C). *Danhatchia australis* (147,326 bp) is similar in size to *S. sinensis* but has a slightly smaller LSC region, while *Chamaegastrodia inverta* (116,220 bp) shows more pronounced reduction in both the LSC and SSC regions but a similar size of the IR relative to that in the two aforementioned species. *Corybas cryptanthus* (69,300 bp) lacks an IR, whereas the IR of both *Rhizanthella gardneri* and *Degranvillea dermaptera* are similarly reduced in size relative *Spiranthes*, *Danhatchia*, and *Chamaegastrodia* (9768 and 11,972 bp, respectively; Fig. 2C). *Degranvillea*, the smallest plastome yet sequenced among subfamily Orchidoideae, has a slightly smaller SSC but a substantially smaller LSC than *Rhizanthella*.

### Phylogenomic analyses

The final plastid gene alignment for Orchidoideae contained 104 taxa with 77,958 aligned positions from 78 genes, 23,711 distinct site patterns, 17.3% missing data or gaps, and 14,485 parsimony-informative characters. Among the four GHOST heterotachy models compared, GTR+FO*H6 had the lowest BIC score (940,961.215; log-likelihood = - 463,254.799; Table 2) and highest BIC weight (wBIC = 1). Overall support was strong for the topology based on this model (Fig. 4). Representatives of orchidoid subtribe Orchidinae Dressler & Dodson ex Reveal were recovered as monophyletic (BS = 100) and sister of *Codonorchis* Lindl., which comprises tribe Codonorchideae P.J. Cribb. This clade together was sister of all remaining ingroup taxa (BS = 100) which comprised two sister clades corresponding to tribes Diurideae Endl. ex Butzin and Cranichideae. Representatives of five subtribes (Acianthinae Schltr., Drakaeinae Schltr., Prasophyllinae Schltr., Rhizanthellinae R.S. Rogers, and Thelymitrinae Lindl. ex Meisn.), which belong to tribe Diurideae, formed a clade (BS = 100) that was sister of subtribes Cranichidinae, Spiranthinae, and Goodyerinae (BS = 100 for the monophyly of all aforementioned clades and for their sister relationships). *Degranvillea dermaptera* was clearly placed within the orchidoid subtribe Spiranthinae, which was strongly supported as monophyletic (BS = 100). More specifically, *Degranvillea* was supported as sister of a clade comprising representatives of the South American genera *Cyclopogon* and *Sauroglossum* (BS = 100). Based on the current sampling, the representatives of the other four leafless species were also strongly supported. *Rhizanthella* (subtribe Rhizanthellinae) was placed as sister of *Microtis* R. Br. (subtribe Prasophyllinae) (BS = 99). *Corybas cryptanthus* was placed within a clade comprising six other species of *Corybas*, and as sister of *Corybas diemenicus* (Lindl.) Rupp (BS = 92). The leafless *Danhatchia australis* was placed as sister of a large clade within subtribe Goodyerinae comprising several genera (BS = 100), and the leafless *Chamaegastrodia inverta* and *C. shikokiana* were placed as sister of a clade comprising three species of *Odontochilus* (BS = 100).

**Fig. 4.**
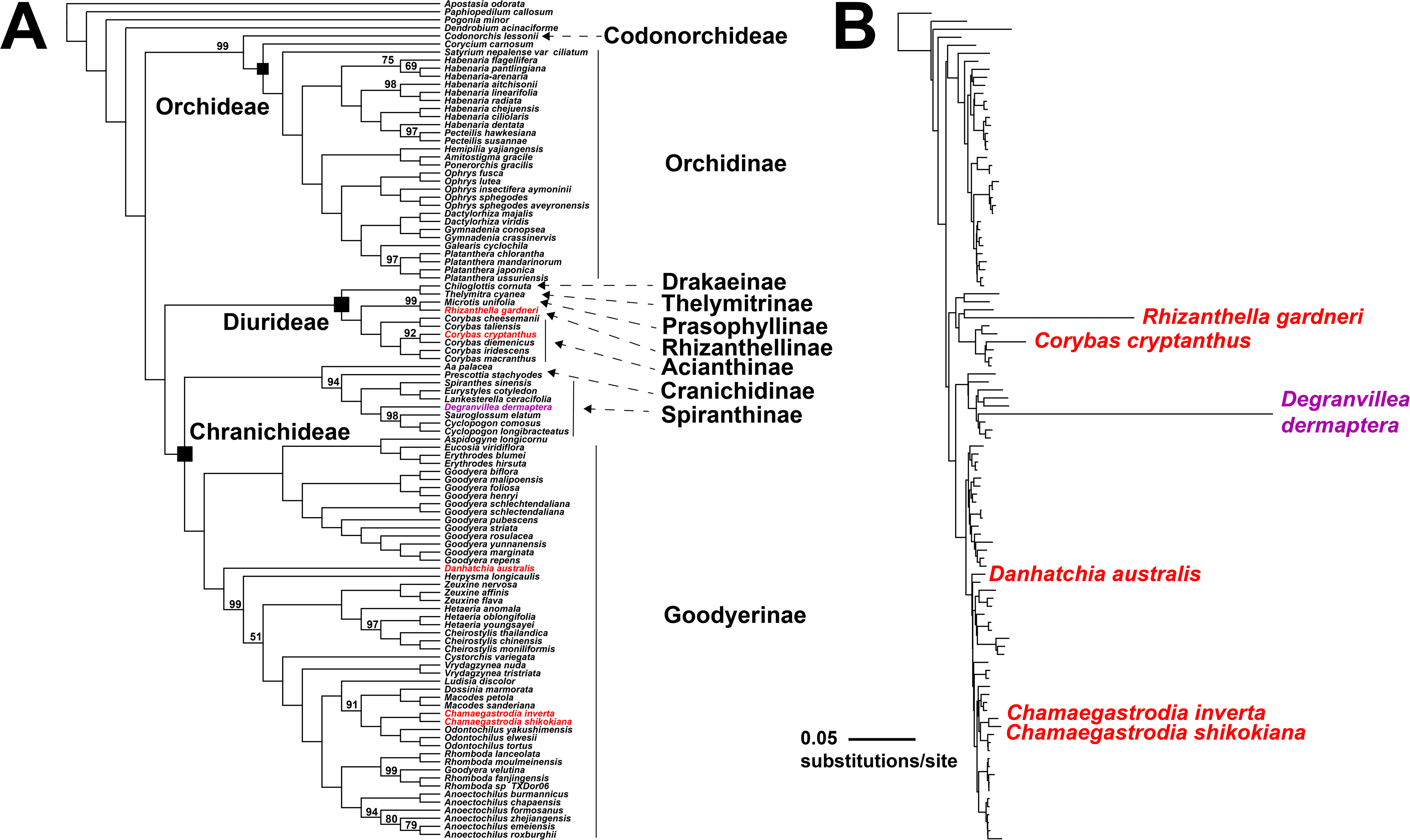
**A.** Cladogram resulting from maximum likelihood phylogenomic analysis among members of the orchid subfamily Orchidoideae based on available plastid genome data (77,958 aligned positions from 78 genes, with 14,485 parsimony-informative characters) under the GHOST heterotachy model in IQTree2 (model = GTR+F0*H6, log-likelihood = -463,254.799, Bayesian Information Criterion = 940,961.215, BIC weight = 1). Numbers above branches are support values from 2,000 ultrafast bootstrap pseudoreplicates in IQTree2; no value indicates 100% support. Black squares indicate three of the four tribes within subfamily Orchidoideae, plus one representative of tribe Codonorchideae (*Codonorchis,* no square shown). Names on the right indicate subtribes. Leafless taxa are colored in red, while the leafless *Degranvillea dermaptera* is colored in magenta. **B**. Phylogram from the same analysis showing estimated branch lengths, with leafless taxa colored as above. Here, branch lengths from the GHOST model are averaged across six different site classes. Scale bar = 0.05 substitutions per site.

**Table 2.** Bayesian Information Criterion (BIC) comparison among phylogenetic GHOST heterotachy models with different numbers of rate classes (K). Delta-BIC is the difference between each BIC value and that of the model with the lowest BIC score. BIC-weight is the relative likelihood of each model. ‘*’ = the best-fit non-heterotachous model identified with ModelFinder.

| Model + rate<br>classes (K) | free parameters | Likelihood | BIC | delta-BIC | BIC-weight |
| --- | --- | --- | --- | --- | --- |
| **GTR+F0\*H (2)** | 427 | -472890.9047 | 950591.505 | 9630.29 | 0 |
| **GTR+F0\*H (4)** | 855 | -466522.586 | 942675.829 | 1714.614 | 0 |
| **GTR+F0\*H (6)** | 1283 | -463254.799 | 940961.215 | 0 | 1 |
| **GTR+F0\*H (8)** | 1455 | -462368.72 | 944010.017 | 3048.802 | 0 |
| **GTR+F+R4 (1)\*** | 219 | -474829.611 | 946440.836 | 5479.621 | 0 |

Coverage depth of the ITS region for *Degranvillea demaptera* was 749.1×. The ITS alignment for Spiranthinae consisted of 317 sequences and 798 aligned positions, with 625 distinct site patterns, 461 parsimony-informative characters, and 17.9% gaps/missing data. Support values were generally strong along the ITS tree backbone (Fig. 5), albeit generally lower than those in the plastome tree. The tree consisted of three principal clades within Spiranthinae: 1) a clade comprising several representative genera including *Eltroplectris* Raf., *Sacoila* Raf., and *Stenorrhynchos* Rich. ex Spreng. (BS = 100); 2) a clade of *Degranvillea* + several genera including *Aulosepalum* Garay*, Dichromanthus* Garay*, Deiregyne* Schltr., *Eurystyles* Wawra, *Hapalorchis* Schltr., *Schiedeella* Schtr., and *Spiranthes* (BS = 95); and 3) a large clade including *Brachystele* Schltr.*, Cyclopogon, Pelexia, Sarcoglottis,* and *Sauroglossum* (BS = 100).

**Fig. 5.**
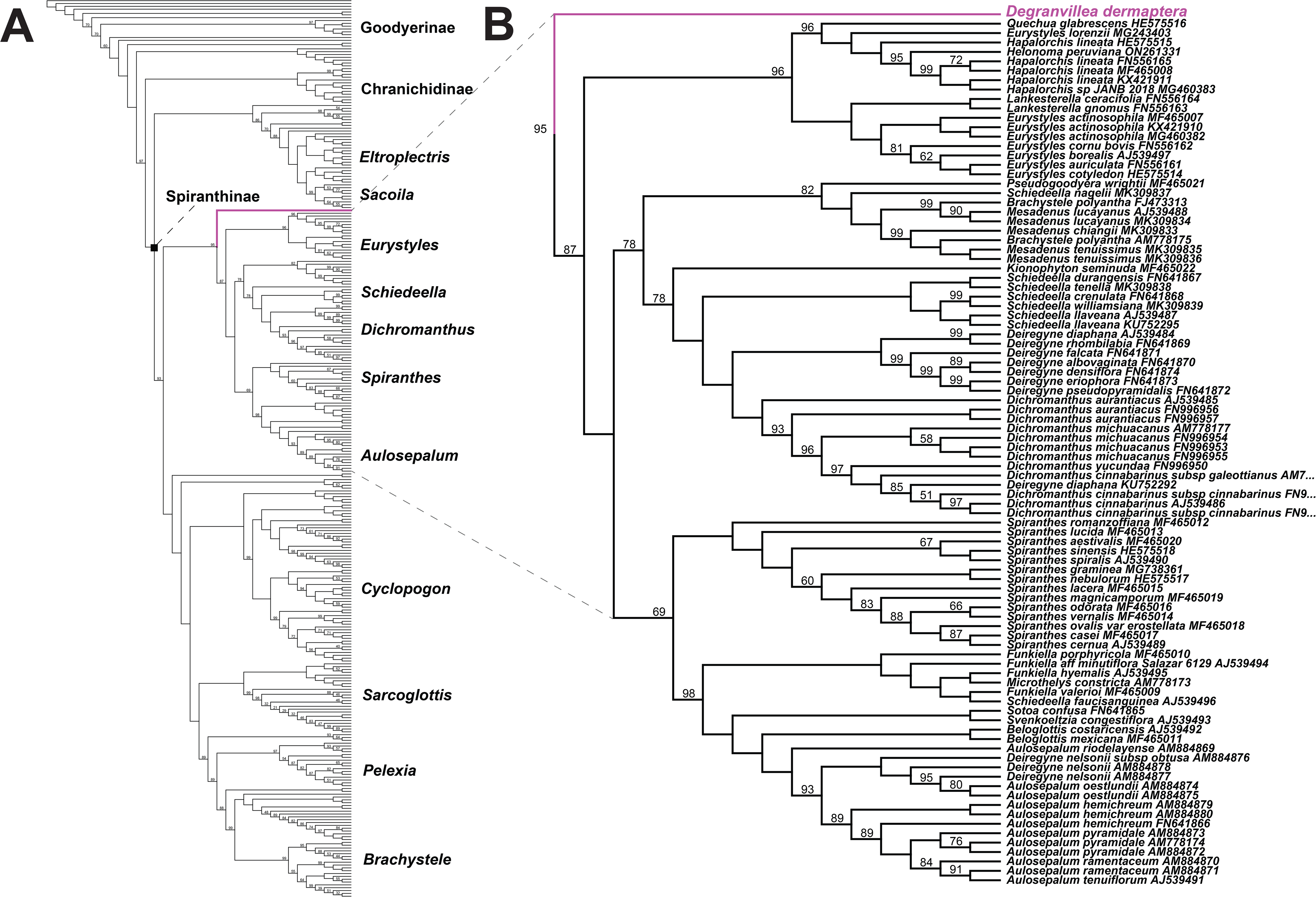
**A.** Cladogram overview resulting from maximum likelihood phylogenetic analysis in IQTree2 among members of the orchid subtribe Spiranthinae based on available nuclear ITS data, with *Degranvillea dermaptera* in magenta (data from Salazar et al., 2018; 317 sequences, 798 aligned positions, 461 parsimony-informative characters, and 17.9% gaps/missing data) (model = TIM3e+I+G4, log-likelihood = -20988.793, Bayesian Information Criterion = 46160.586). Select genera are listed. Black square indicates subtribe Spiranthinae. **B.** Close-up of the clade containing *Degranvillea dermaptera* (in magenta). Numbers above branches are bootstrap support values from 2,000 ultra-fast pseudoreplicates in IQTree2, with blank values indicating 100% support.

Divergence time estimates for the initial diversification of Orchidoideae, based on our plastid analyses, were 33.89-43.37 and 43.61-55.60 mya (95% confidence intervals) for our minimum/maximum calibration points, respectively (Appendix S4). Estimates for leafless species and their closest sampled leafy relatives placed *Degranvillea* as diverging from the common ancestor of *Cyclopogon* and *Sauroglossum* at 17.37-24.04 and 18.92-27.03 mya. Estimates for the *Rhizanthella-Microtis* divergence ranged from 28.96-39.85 to 34.58-48.14 mya, and *Corybas cryptanthus-C. diemenicus* from 6.16-10.13 to 7.80-12.75 mya. *Chamaegastrodia inverta* and *shikokiana* were estimated to have diverged between 3.99-6.37 and 6.01-9.71 mya, while the most recent common ancestors of *Chamaegastrodia* and *Odontochilus* were estimated to have diverged between 5.20-7.39 and 6.01-9.71 mya. Finally, *Danhatchia australis* was estimated to have diverged between 13.25-15.00 and 18.00-20.01 mya from the ancestor of a large sister clade of several genera within Goodyerinae. Analyses based on ITS data (Appendix S5) displayed slightly older but similar divergence times for *Degranvillea* compared to those from plastid data: 21.85-34.33 mya (15 my calibration) and 23.11-34.08 mya (20 my calibration).

### Genome evolutionary analyses

Analyses within the RELAX framework of HyPhy, with representatives of all five leafless genera specified as test branches, revealed significantly relaxed selection pressure for seven genes (*rpl16, rpl23, rpl36, rps2, rps3, rps4, rps11*), and intensified selection for one other (*rpl20*) (Table 3). A second analysis, considering only *Rhizanthella* and *Degranvillea* as test branches, also revealed significantly relaxed pressure for seven genes (*rpl14, rpl16, rps2, rps4, rps11, rps14,* and *rps16*) and intensified selection for one (*rps3*), but notably these were not all the same genes as in the first analysis (Table 3). Also of note, the latter genes with evidence of significantly relaxed selection are all situated either partially or fully within the SSC region of *D. dermaptera* (corresponding to the LSC in other orchid taxa), and all are in proximity to the boundaries of IR_B_ (*rpl16, rpl14, rps11*) or IR_A_ (*rps4, rps14, rps2, rps16*) (Fig. 3).

**Table 3.** Results of RELAX analyses in HyPhy. ‘K’ is the selection intensity parameter, ‘p-value’ indicates the significance of the likelihood ratio test between the null and alternative models, and ‘Selection’ indicates relaxation or intensification of selection pressure. In Analysis 1, five leafless taxa were specified as test branches (*Chamaegastrodia inverta, Corybas cryptanthus, Danhatchia australis, Degranvillea dermaptera,* and *Rhizanthella gardneri*); in Analysis 2, test branches were restricted to *Degranvillea dermaptera* and *Rhizanthella gardneri*.

| Gene | Analysis 1 (5 leafless spp.) |  |  | Analysis 2 (*Degranvillea*, *Rhizanthella*) |  |  |
| --- | --- | --- | --- | --- | --- | --- |
|  | K | p-value | Selection | K | p-value | Selection |
| *accD* | 1.18 | 0.452 | intensification | 10.63 | 0.115 | intensification |
| *clpP* | 0.12 | 0.707 | relaxation | 0.93 | 0.802 | relaxation |
| *rpl2* | 0.84 | 0.413 | relaxation | 0.74 | 0.2 | relaxation |
| *rpl14* | 0.56 | 0.2 | relaxation | 0.5 | 0.013 | **relaxation** |
| *rpl16* | 0.62 | 0.004 | **relaxation** | 0.57 | 0.022 | **relaxation** |
| *rpl20* | 3.36 | <0.001 | **intensification** | 0.8 | 0.477 | relaxation |
| *rpl23* | 0.012 | 0.01 | **relaxation** | 0.2 | 0.058 | relaxation |
| *rpl36* | 0.17 | 0.025 | **relaxation** | 0.18 | 0.211 | relaxation |
| *rps2* | 0.68 | 0.016 | **relaxation** | 0.68 | 0.028 | **relaxation** |
| *rps3* | 0.19 | 0.003 | **relaxation** | 44.85 | 0.024 | **intensification** |
| *rps4* | 0.49 | <0.001 | **relaxation** | 0.56 | 0.005 | **relaxation** |
| *rps8* | 0.9 | 0.526 | relaxation | 0.85 | 0.356 | relaxation |
| *rps11* | 0.44 | <0.001 | **relaxation** | 0.42 | <0.001 | **relaxation** |
| *rps12* | 0.59 | 0.063 | relaxation | 0.61 | 0.099 | relaxation |
| *rps14* | 0.05 | 0.084 | relaxation | 0.46 | 0.233 | **relaxation** |
| *rps16* | 0.36 | 0.697 | relaxation | 0.46 | 0.233 | **relaxation** |
| *rps18* | 0.66 | 0.23 | relaxation | 0.001 | 0.001 | relaxation |
| *rps19* | 0 | 0.653 | relaxation | 18.43 | 0.396 | intensification |
| *ycf1* | 1.1 | 0.325 | intensification | 50 | 0.533 | intensification |
| *ycf2* | 0.87 | 0.825 | relaxation | 1.28 | 0.276 | intensification |

Analysis with CoEvol revealed several significant correlations among plastome features (Fig. 6). Both the synonymous (*dS*) and nonsynonymous substitution rates (*dN*) had significant positive correlations (posterior probability, or PP >0.95) with root to tip patristic (RTT) distance, double cut and join distance (DCJ, or inversion distance), and microsatellite density, while they were both negatively correlated (PP <0.05) with plastome length, the number of CDS (CDS) and tRNAs (tRNA), GC content (GC), and total shared intron length. *dN* was also positively correlated with *dS,* and negatively correlated with the number of perfect repeats >20 bp. Plastome length was positively correlated with CDS, tRNA, GC content, repeats > 20 bp, and shared intron length. It was negatively correlated with RTT and DCJ distances. CDS was positively correlated with tRNA, GC, repeats >20 bp, and intron length. It was negatively correlated with RTT and DCJ distances. tRNA was correlated with RTT distance (+), intron length (+), and RTT distance (−). RTT distance was correlated with intron length (−), microsatellite density (+), and GC content (−). Lastly, GC content was correlated with microsatellite density (−) and repeats >20 bp were correlated with intron length (+).

**Fig. 6.**
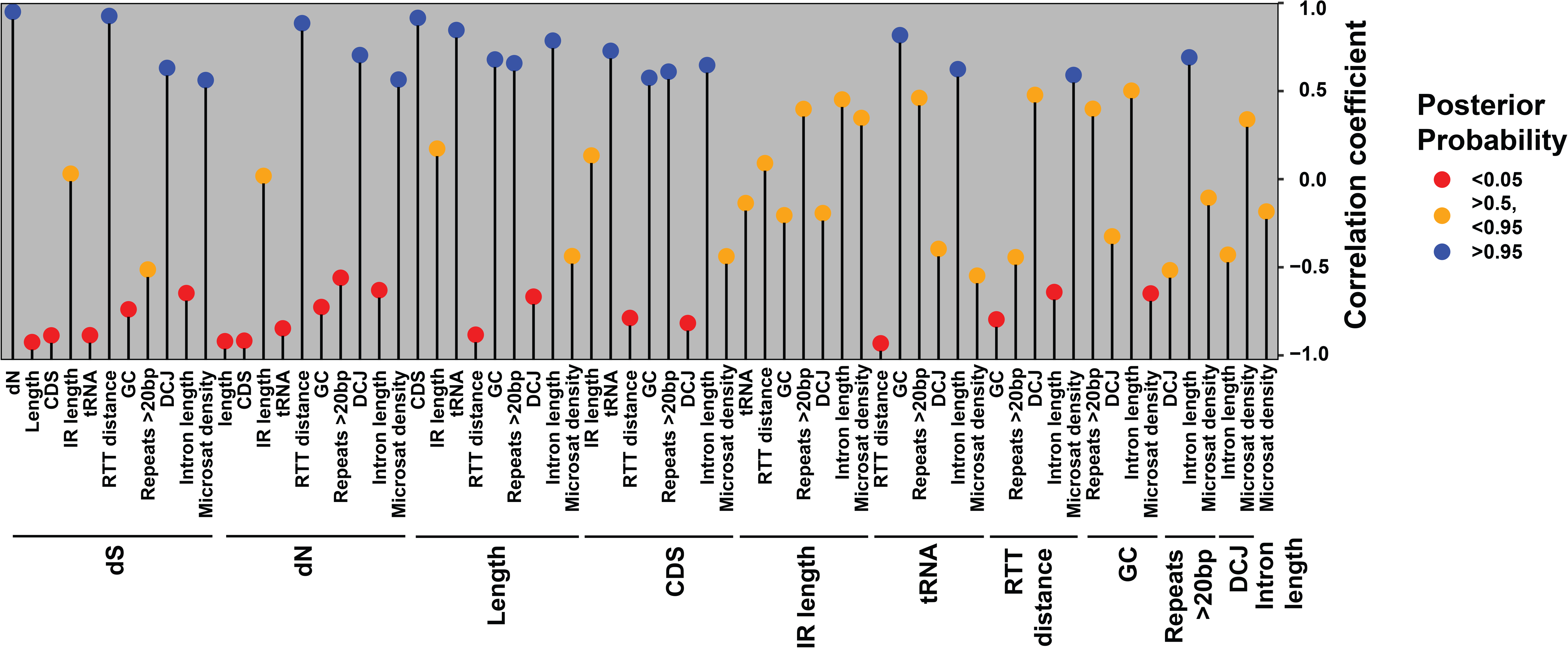
Results of CoEvol analysis, showing all pairwise comparisons among two substitutional features (*dN*, *dS*) and ten other features of the plastome. Comparisons in boldface received posterior probabilities >0.95 or < 0.05, suggesting significant negative or positive correlations, and their correlation coefficients are shown in red or blue, respectively. Comparisons with probabilities >0.05 or <0.95 are in orange. ‘*dS*’ = the proportion of synonymous substitutions per synonymous site, ‘*dN*’ = the proportion of nonsynonymous substitutions per nonsynonymous site, ‘Length’ = plastome length, ‘IR length’ is the length of the Inverted Repeat region, ‘CDS’ = the number of putatively functional protein coding genes, ‘tRNA’ = the number of putatively functional transfer RNA genes, ‘RTT distance’ = root-to-tip patristic distances based on the GHOST heterotachy model (GTR+F0*H6), ‘GC’ = GC content, and ‘Repeats >20 bp’ = the number of perfect repeats ≥20 bp in length, ‘DCJ’ = double cut and join distance (inversions), ‘Intron length’ is the sum in bp of the four introns retained in all species, and ‘Microsat density’ is the number of detected miscrosatellites divided by total plastome length (all metrics consider plastomes with one IR copy removed).

## DISCUSSION

We sequenced and assembled the smallest plastome known thus far among the orchidoid orchids, a clade dominated by a terrestrial habit that comprises nearly 3800 species (Pridgeon et al., 2001; 2003; Chase et al., 2015; Salazar et al., 2018). We revealed a highly reduced plastome in total length and functional gene content, which lacks all photosynthesis-related genes, most transfer RNA genes, and several “housekeeping” genes, but retains all five “core nonbioenergetic” genes (*sensu* Graham et al., 2017). We found evidence for structural rearrangements, elevated substitution rates, relaxed selection on several retained genes, and correlations between substitutional parameters and other “traits” of the plastome among leafless orchidoids. We found strong evidence based on plastomes and nuclear ITS for the placement of *Degranvillea* within the subtribe Spiranthinae, corroborating earlier hypotheses based on morphological affinity (Determann, 1985; Chase et al., 2015; Salazar et al., 2018). The study advances knowledge on phylogenetic relationships and the trajectory of evolution within subfamily Orchidoideae, and heterotrophic plants more broadly, and demonstrates the power of herbarium collections in comparative phylogenomics.

### Genome structure

The path of plastome evolution in *D. dermaptera* appears to follow several previous models of genome degradation in heterotrophic plants (Wicke, 2011; Barrett and Davis, 2012; Barrett et al., 2014; Wicke et al., 2016; Graham et al., 2017; Wicke and Naumann, 2018). A comparison among sequenced leafless orchids and leafy relatives (Fig. 3) reveals loss of the entire NADPH complex (*ndh*) in *Degranvillea* and nearly all leafless taxa, including both partial and full mycoheterotrophs. In fact, only a few fully mycoheterotrophic species retain any *ndh* genes (e.g. *Aphyllorchis, Corallorhiza, Diplandorchis*). These tend to be the first genes lost in the transition to heterotrophy, but *ndh* loss is not restricted to heterotrophs, as their function in electron recycling is thought to be dispensable or redundant in some photosynthetic lineages (Wicke et al., 2011; Peltier et al. 2016). In fact, evidence for *ndh* gene loss is found among at least some leafy, photosynthetic species in all five orchid subfamilies (e.g. Kim et al., 2020). We detected severely truncated pseudogenes in *Degranvillea* for two *ndh* genes (*ndhB* and *ndhF*), which in most species are fully or partially embedded in the IR. The IR tends to display slower rates of sequence evolution, generally speaking, than either Single Copy region (Wolfe et al., 1987; Perry and Wolfe, 2002; Mower and Vickrey, 2018; Wang et al., 2022), which could explain why fragments of these genes have persisted. Wicke et al. (2016) suggested that proximity to more highly conserved genes or gene regions essential for core plastid function may result in a “ripple effect,” and thus nearby, otherwise dispensable genes may be protected from wholescale loss due to genetic hitchhiking (in terms of functional constraint).

*Degranvillea* retains no functional or even pseudogenized copies of photosynthesis-related genes. These include, for example, genes encoding Photosystems I and II (*psa, psb*), Cytochrome (*pet*), Photosystem Assembly factors (*pafI* and *pafII*, formerly referred to as *ycf3* and *ycf4*), and the RuBisCO Large subunit (*rbcL*). Other genes indirectly involved in photosynthesis have been lost in *Degranvillea*, including those for the Plastid-Encoded RNA Polymerase (*rpo*) and ATP Synthase (*atp*). It is intuitive as to why the *rpo* complex would be lost, either following or concomitant with the loss of e.g. *psa* and *psb* genes, as this polymerase transcribes most of these photosynthesis-related genes (Liere et al. 2011). The loss of *atp* genes, however, is more intriguing. ATP Synthase plays a critical role in photosynthesis by phosphorylating ADP to ATP, but it also maintains a proton gradient across the chloroplast thylakoid membrane which in turn is essential for proper function of a Twin-Arginine Translocator complex that transports folded proteins across the thylakoid membrane (Kamikawa et al. 2015; New et al., 2018; discussed in Barrett et al., 2019; Cheuk and Meyer, 2021). The loss of ATP Synthase function, and hypothesized lack of a proton gradient required for the essential process of protein translocation across the thylakoid membrane, may mean that plants lacking a functional *atp* complex could either harbor dysfunctional chloroplasts, or that they have lost chloroplasts altogether. Clearly this is an example of how research in heterotrophic organellar biology has been outpaced by genomic sequencing. It remains to be determined whether late stage heterotrophs even have chloroplasts as opposed to other types of plastids. There is a clear need for more cellular research in these plants including detailed microscopy to elucidate the dynamics of basic plastid biology.

Several “housekeeping” genes, e.g., Ribosomal Protein genes and genes encoding transfer RNAs (*rpl, rps, trn*) involved in plastid gene translation, are missing altogether, suggesting *Degranvillea* is in the fifth, and penultimate stage of degradation more or less (*sensu* Barrett and Davis, 2012). *Degranvillea* also retains what Graham et al. (2017) define as the five core nonbioenergetic genes (*trnE, accD, clpP, ycf1, and ycf2*). While the functions of these genes are explained elsewhere in detail, Graham et al. (2017) note that among them, *accD* (encoding a subunit of acetyl-CoA carboxylase involved in fatty acid biosynthesis) and *trnE* (encoding glutamyl transfer RNA but also functioning in tetrapyrrole biosynthesis) tend to be “sticky,” meaning that they are often among the last remaining functional genes to be retained in the plastome, even among the most reduced species sequenced to date. While we consider *Degranvillea* to be in the late stages of degradation, it is by far not the smallest plastome among heterotrophic plants. Among orchids, *Pogoniopsis schenckii* Cogn., *Didymoplexis pallens* Griff., two species of *Epipogium* J.F. Gmel. ex Borkh., and several species of *Gastrodia* R. Br. Have smaller plastomes and similar gene content, with loss or severe reduction of the IR explaining the smaller size in some of these species (Fig. 3).

Of note is the loss of *matK* in *Degranvillea*, which is demonstrated to function in splicing of Group IIA introns (Vogel et al.,1999; Bonen and Vogel, 2001; Zoschke et al., 2010), despite retention of those introns in *3’-rps12* and *rpl2* (Appendix S6). Several scenarios can be hypothesized for the loss of *matK*: 1) these intron-containing genes are self-splicing; 2) they are spliced by some other mechanism and not the product of *matK*; 3) a functional *matK* copy has been transferred to another genomic compartment and its product is imported into the plastome; 4) *matK* has been lost altogether in some lineages (e.g. *Degranvillea*) and thus genes with Group IIA introns are transcribed and translated, somehow remaining functional; 5) these genes are transcribed but result in non-functional products; or 6) they are not transcribed at all, and may be effectively considered pseudogenes (Zoschke et al., 2010; Delannoy et al., 2011; Naumann et al., 2016; Graham et al., 2017). The situation in the epidendroid orchid *Pogoniopsis schenckii* (Klimpert et al., 2022), which has lost both *matK* and all Group IIA introns (as might be expected given the loss of *matK*), may represent what the future holds for *Degranvillea*, with the latter representing an intermediate stage in plastome reduction, considering gene-gene product interactions. A similar situation is observed among several species of *Gastrodia* and the related *Didymoplexis pallens* which have lost *matK* (independently within tribe Gastrodieae Lindl.) but retain the same Group IIA introns as *Degranvillea* (Wen et al., 2022).

The plastome of *Degranvillea* has experienced major structural rearrangements, in line with what might be expected in a highly modified plastome (e.g. Wicke et al., 2016). The IR region is much reduced, but comparable to that in similar-sized plastomes (e.g. some *Gastrodia*, Wen et al., 2022; *Rhizanthella*, Delannoy et al., 2011; *Epipogium aphyllum* Sw., Schelkunov et al., 2015). Interestingly, the ancestral LSC region in *Degranvillea* is smaller than the region corresponding to the SSC (Figs. 2, 4), largely due to losses of genes and gene regions in the ancestral LSC and contraction of the IR at the SSC boundaries (Fig. 4). In terms of gene order, there are two major rearrangements relative to the photosynthetic *Spiranthes sinensis*, involving inversions of *accD* (LSC) and *ycf1* (SSC; Fig. 2B). Notably, there is also evidence of palindromic repeats near the hypothesized breakpoints of the *accD* inversion (Appendix S3), which are hypothesized to facilitate such rearrangements (e.g. inversions, deletions, translocations) as have been observed in other taxa such as *Asarum* L. (family Aristolochiaceae) and several lineages within order Asterales Link (Knox, 2014; Sinn et al., 2015; Knox, 2022). Some heterotrophic taxa with highly reduced plastomes maintain a structure that is colinear with leafy, photosynthetic relatives (e.g. *Sciaphila* Blume, Lam et al., 2015), while others do not, and the reasons for these seemingly idiosyncratic evolutionary trajectories is unknown. The presence of an IR has been hypothesized to promote stability in plastome synteny, but there is evidence both for and against this hypothesis. Some lineages with a seemingly “standard” IR experience extensive rearrangements, while others with highly reduced or missing IR regions do not (reviewed in Wang et al., 2022).

### Phylogenomic analyses

The results of our phylogenetic analyses clearly placed *Degranvillea* within the orchidoid subtribe Spiranthinae, and thus far to our knowledge, *Degranvillea* represents the sole, fully mycoheterotrophic member of this species-rich clade (Figs 5, 6). These findings corroborate earlier hypotheses based on morphological similarity to members of Spiranthinae (Determann, 1985; Salazar, 2018), but the lack of morphological features in the highly reduced *Degranvillea* has made taxonomic placement a challenge. While our analysis based on plastid genes allowed phylogenetic placement with high support at the subtribal level, too few representatives of Spiranthinae have been sequenced to date to allow species- or genus-level resolution of the placement of *Degranvillea*. Fortunately, the availability of nearly complete generic-level sampling of nuclear ITS sequences for this subtribe (Salazar et al., 2018) allowed a more fine-scale phylogenetic analysis (*viz*. taxon sampling but not character sampling), revealing that *Degranvillea* occupies a position as sister of a species-rich clade of predominantly Central and South American genera (e.g. *Aulosepalum, Dichromanthus, Deiregyne, Erystyles, Funkiella, Hapalorchis, Mesadenus, Schiedeella*), in addition to *Spiranthes*, which has diversified into North America, Eurasia, and Australia (Dueck et al., 2014; Givnish et al., 2015; Serna-Sanchez et al., 2021; Perez-Escobar et al., 2023). Salazar et al. (2018) also included matching taxon sampling for the plastid *matK* and *trnL-F* regions, but both of these regions have been lost in *Degranvillea*, thus limiting our analysis to a single locus in ITS. It is curious in our analysis that *Degranvillea*, a leafless, monotypic genus endemic to French Guiana, is sister of a clade comprising several hundred species (Fig. 6). While many leafless orchid taxa are represented by multiple species (e.g. *Aphyllorchis, Corallorhiza, Didymoplexis, Diodium, Gastrodia, Hexalectris, Lecanorchis*), others including *Degranvillea* are not (Merckx et al., 2013a). Some have suggested that leaflessness, full mycoheterotrophy, and photosynthetic loss may represent an evolutionary “dead end,” but the evidence for this is equivocal (Bidartondo, 2005; Waterman et al., 2008; Guo et al., 2019). Perhaps the question is unanswerable due to the exceedingly poor fossil record of orchids in general, and mycoheterotrophs more specifically, for which no fossils exist due to their typically small stature, low population densities, and reduction in vegetative tissues.

Spiranthinae is almost entirely terrestrial. However, after the divergence of mycoheterotrophic *Degranvillea*, the next branching event within Spiranthinae includes the divergence of *Spiranthes* and many Mexican and Central American relatives, and the evolution of the only lithophytic (*Helonoma* and *Quechua*) and epiphytic (*Eurystyles*, *Lankesterella,* and *Pseudoeurystyles*) genera within the subtribe. It is likely that the ancestors of *Degranvillea*, *Eurystyles*, *Helonoma*, *Lankesterella*, *Quechua*, and *Pseudoeurystyles*, and the clade involving *Spiranthes* and its most closely related genera were terrestrial and photosynthetic, however it is intriguing that both the diversification of habit and the evolution of mycoheterotrophy are clustered in this small section of the wider Spiranthinae. The question of “why” immediately arises, and future research should be directed towards addressing this topic with increased taxon sampling and the application of ancestral state and diversification analyses. In this context, the biogeographic endemism of *Degranvillea* to the Guiana Shield is also intriguing. The biogeographic analysis of Salazar et a. (2018) did not code the Guiana Shield as a distinct area, appearing to include it within “Eastern South America.” This was probably done because they were unable to sample the Spiranthinae genera that are endemic to the Guiana Shield, and the species they did include from this region also occur elsewhere in tropical America. Based on our phylogenetic analysis of *Degranvillea*, combined with those of Salazar et al. (2018, 2022), we hypothesize that the Guiana Shield is a possibly a sink for Spiranthinae, where lineages migrate to but do not migrate from. While the Guiana Shield apparently has not promoted major speciation events in this clade, it may have promoted the evolution of unusual habits relative to the rest of the subtribe, including the mycoheterotrophic *Degranvillea*, the (likely) carnivorous *Aracamunia* Carnevali & I. Ramírez (Steyermark and Holst, 1989), and the epiphytic *Helonoma* Garay.

Our divergence time analyses, considering our conservative approach using both minimum and maximum ages for the fossil *Meliorchis* in Goodyerinae and both the plastid and ITS analyses, placed the divergence of *Degranvillea* from its sister clade within Spiranthinae at some time between 17.37 and 34.43 mya (Appendices S5, S6). Thus, it is possible that it has taken at most 35 my for the plastome of this species to experience a nearly 70% reduction in overall length and a 75% reduction in functional gene content compared to that in leafy relatives. While it is tempting to make comparisons of divergence times among the five orchidoid genera included in this analysis, we refrain from doing so due to the dearth of sequenced plastomes and the need for—at a minimum—genus-level sampling. That said, there are several meaningful analyses of leafless/leafy divergence times available for comparison. At one extreme, the endoparasitic family Rafflesiaceae was estimated to have diverged from the closest leafy relative ca. 101 mya (confidence interval = 96-107 mya; Pelser et al., 2019), meaning at most it would have taken ca. 100 my for complete loss of the plastome in this clade of endoparasites (Molina et al., 2014). The ancestor of the epidendroid orchid tribe Gastrodiae is estimated to have diverged from the leafy tribe Nervilieae Dressler ca. 36 mya. Plastomes of sequenced Gastrodiae range from 51,241 bp in *Didymoplexis pallens* to 29,626 bp in *Gastrodia peichatieniana* S.S. Ying, with similar gene content to *Degranvillea* and *Rhizanthella* among the orchidoids (Wen et al., 2022).

By contrast, the leafless epidendroid orchid genus *Corallorhiza* Gagnebin and closest leafy relative *Oreorchis* Lindl. were estimated to have diverged ca. 10.4 mya (Barrett et al., 2018). Plastome sizes range from being similar among leafless and leafy species between the two genera (151,506 bp in the leafless, partially mycoheterotrophic *C*. *maculata* var. *mexicana* (Lindl.) Freudenstein and 158,654 bp in the leafy *O. angustata* L.O. Williams ex N. Pearce & P.J. Cribb to 124,420 bp in the leafless, fully mycoheterotrophic *C. involuta* Greenman. A remarkable example is that of the monocot family Thismiaceae J. Agardh. This family comprises five genera and is estimated to have diverged from the leafy sister family, Taccaceae Dumort., ca. 11.51 mya based on plastome data (Givnish et al., 2018; Shepeleva et al., 2020). If this estimate is accurate, then in under ca. 15 my members of this family have lost the vast majority of the plastome (size range: *Haplothismia exannulata* Airy Shaw = 22,294 bp, *Thismia hexagona* Dančák, Hroneš, Kobrlová & Sochor = 5904 bp; Yudina et al., 2021; Garrett et al., 2022). However, earlier divergence time estimates place the stem age of Thismiaceae up to 100 mya (Merckx and Smets, 2014; Hertweck et al., 2015; Merckx et al., 2017; reviewed in Garrett et al., 2022), highlighting the challenges in divergence time estimation in heterotrophic lineages with highly reduced plastomes and massively accelerated substitution rates. While data and comparisons are still too few to conduct a global analysis of the relationship between leafy-leafless divergence times and the degree of plastome degradation, efforts to fill sampling gaps necessary to do so should be a future priority. Though extinction may be a confounding factor, such a comparison would result in a relative timeframe of plastome degradation, and further would allow more fine-scale investigation of the factors (molecular, physiological, environmental, etc.) that drive the rate of genome modification across different lineages.

### Genome evolutionary analyses

We found evidence of relaxed selection in several retained “housekeeping” genes, all of which were Ribosomal Protein subunits (Table 2). Interestingly, several of these genes are located near the IR boundaries in *Degranvillea*, suggesting the possibility that “release” from the substitutional constraints of IR occupancy may have led to relaxed selection on these genes. Wang et al., (2022) revealed that overall substitution rates in IR-lacking species were higher than closely related IR-containing species in some, but not all comparisons. Genes in the IR can experience up to ∼4× lower substitution rates than their counterparts outside the IR (Wolfe et al., 1987; Perry and Wolfe, 2002; Mower and Vickrey, 2018; Wang et al., 2022). When considering only *Degranvillea* and *Rhizanthella* as test branches, we found evidence of intensified selection in *rps3*, which is situated within the IR in *Degranvillea* but lies at the LSC-IR_B_ boundary in *Rhizanthella* (Fig. 2).

In heterotrophic plants both nonsynonymous (*dN*) and synonymous (*dS*) substitution rates may experience increases simultaneously, thus making it challenging to rely on metrics of relaxed selection such as *dN*/*dS* ratios (e.g. Bromham et al., 2013). Indeed, the strongest correlation in our CoEvol analysis was between *dN* and *dS*, suggesting overall elevated substitution rates in heterotrophic orchidoid lineages. These two substitutional parameters, in turn, were correlated with most “traits” of the plastome including total plastome length (−), branch lengths (root-to-tip patristic distance, +), inversions (+), GC content (−), the numbers of CDS and tRNAs (−), repeat content and density (+), and retained intron length (−) (Fig. 6). These findings corroborate earlier comparative studies in which a suite of correlated changes characterize plastomes in heterotrophs [e.g. Orobanchaceae Vent., Wicke et al., 2016; *Corallorhiza*, *Gastrodia*, and *Hexalectris* Raf. (Orchidaceae), Barrett et al., 2018; 2019; Wen et al., 2022; Thismiaceae, Garrett et al., 2023; Balanophoraceae Rich., Kim et al., 2023].

## CONCLUSIONS

We have sequenced and analyzed the most reduced plastid genome to date among the orchid subfamily Orchidoideae, which comprises ∼15% of orchid biodiversity and contains numerous mycoheterotrophic species. Our study underscores the crucial role of herbarium collections in genomic research for rare species of conservation concern, and fills an important knowledge gap on phylogenetic relationships and evolutionary dynamics of plastid genome evolution within this clade. In particular, we found strong support and corroborated earlier hypotheses for the phylogenetic placement of *Degranvillea dermaptera*, a poorly studied and rarely collected species, within the orchid subtribe Spiranthinae. However, even with the current high rate and continued interest in plastid genome sequencing, there are numerous sampling gaps (including heterotrophic lineages) that hinder our ability to conduct comprehensive analyses comparing the rate and degree of genome modification among independently derived instances of heterotrophy in plants. Increased sampling of heterotrophic and photosynthetic lineages will allow researchers to identify the underlying similarities and idiosyncrasies of plastome evolution, including what drives the rate of evolutionary change among lineages.

## AUTHOR CONTRIBUTIONS

C.F.B. conceived the study, performed DNA extractions, analyzed data, and led the writing of the manuscript. M.P. collected herbarium material and provided the specimen image of *Degranvillea*. C.W.C. assessed DNA quality and prepared sequencing libraries. All authors helped draft and revise the final manuscript.

## ACKNOWLEDGEMENTS

Funding was provided by the US National Science Foundation CAREER award DEB 2044259 to CFB. The specimen of *Degranvillea* was digitized with support from NSF award DBI 1802034 to MP. We acknowledge the historical custodians of what is now French Guiana, where specimens making this work possible were collected, including the Apalaï, Kali’na, Lokono, Palikur, Teko, Wayana, and Wayãpi. We further acknowledge that this work was completed on land that includes ancestral territories of the Shawnee, Lenape (Delaware), Cherokee, Haudenosaunee (Seneca, Cayuga, Onondaga, Oneida, Mohawk, and Tuscarora), and many other Indigenous Peoples. The staff at the New York Botanical Garden William and Lynda Steere Herbarium granted permission to collect material. We are grateful to Scott Mori (1941-2020) for collection of the specimen which made this study possible; we also thank him and Jean-Jacques de Granville (1943-2022, the namesake of *Degranvillea*) for their advocacy in the formation of the Guiana Amazonian Park. Jun Fan, Ryan Percifield, and Donald Primerano provided sequencing support and discussion. We thank Noah Adkins, Damian Disbrow, Lauren Kosslow, Sam Skibicki, and Hana Thixton-Nolan for helpful discussion. We thank the WVU Genomics Core Facility for support provided to help make this publication possible and CTSI Grant no. U54 GM104942 which in turn provides financial support to the WVU Core Facility. We further acknowledge WV-INBRE, a COBRE ACCORD grant, and a West Virginia Clinical and Translational Science Institute (WV-CTSI) grant in supporting the Marshall University Genomics Core.

## DATA AVAILABILITY STATEMENT

The plastid genome of *Degranvillea dermaptera* was deposited to NCBI GenBank under accession number XXXXXXX. Raw data in FASTQ format were submitted to the NCBI Sequence Read Archive (BioProject XXXXXXX). Plastid and nuclear ITS alignments are provided via Zenodo (https://zenodo.org/badge/DOI/10.5281/zenodo.10070346).

## SUPPORTING INFORMATION

**Appendix S1.** Analysis of post-mortem DNA substitutions with MapDamage. **A.-D.** Frequencies of each base outside and inside the read (the open grey box corresponds to the read). **E.-F.** Position-specific substitutions from the 5’ (E) and the 3’ ends (F). Red = C to T substitutions, blue = G to A substitutions, gray = all other substitutions. **G.** Empirical frequencies of base misincorporation (lines) and simulated posterior intervals (points and error bars) from the fitted model.

**Appendix S2**. Inverted Repeat boundaries among the leafy *Spiranthes sinensis* and four leafless taxa (not to scale). *Corybas cryptanthus* is not included due to lacking an IR. LSC = Large Single Copy Region, SSC = Small Single Copy Region, and IR = Inverted Repeat. Numbers in blue indicate the length of each region in base pairs. Arrows and numbers near IR-Single Copy junctions indicate the distance from the start/end of a gene to the junction. ‘JLB’ = LSC-IR_B_ junction, ‘JSB’ = IR_B_-SSC junction, ‘JSA’ = SSC-IR_A_ junction, and ‘JLA’ = IR_A_-LSC junction. The SSC-IR_B_-LSC region of *Degranvillea* was manually reversed to reflect synteny among the regions, since technically the region syntenically corresponding to the LSC in other orchids is shorter than the other Single Copy region within the *D. dermaptera* genome.

**Appendix S3**. Analysis of detected palindromic repeats (below) and the structures of three notable repeats that occur near the putative *accD* inversion breakpoints (above). ‘dG’ = Gibbs Free Energy (Joules).

**Appendix S4**. Least-squares divergence time estimates from plastid gene data. Blue bars and numbers above branches indicate the 95% confidence limits based on 1,000 bootstraps. Leafless taxa are shown as red branches, with *Degranvillea* in magenta. Scale bar = millions of years ago.

**Appendix S5**. Least-squares divergence time estimates from ITS data. Blue bars and numbers above branches indicate the 95% confidence limits based on 1,000 bootstraps. Leafless taxa are shown as red branches, with *Degranvillea* in magenta. Scale bar = millions of years ago.

**Appendix S6**. Intron lengths (bp) for all intron-bearing CDS and tRNA genes among five leafless orchidoid species and the leafy *Spiranthes sinensis*. Blank entries represent physical gene losses. ‘*’ indicates a Group IIA intron.

## REFERENCES

Bankevich, A., S. Nurk, D. Antipov, A. A. Gurevich, M. Dvorkin, A. S. Kulikov, V. M. Lesin, et al. 2012. SPAdes: A new genome assembly algorithm and its applications to single-cell sequencing. Journal of Computational Biology 19: 455–477.

Barrett, C. F., and J. I. Davis. 2012. The plastid genome of the mycoheterotrophic *Corallorhiza striata* (Orchidaceae) is in the relatively early stages of degradation. American Journal of Botany 99: 1513–1523.

Barrett, C. F., J. V. Freudenstein, J. Li, D. R. Mayfield-Jones, L. Perez, J. C. Pires, and C. Santos. 2014. Investigating the path of plastid genome degradation in an early-transitional clade of heterotrophic orchids, and implications for heterotrophic angiosperms. Molecular Biology and Evolution 31: 3095–3112.

Barrett, C. F., S. Wicke, and C. Sass. 2018. Dense infraspecific sampling reveals rapid and independent trajectories of plastome degradation in a heterotrophic orchid complex. The New Phytologist 218: 1192–1204.

Barrett, C. F., B. T. Sinn, and A. H. Kennedy. 2019. Unprecedented parallel photosynthetic losses in a heterotrophic orchid genus. Molecular Biology and Evolution 36: 1884–1901.

Beier, S., T. Thiel, T. Münch, U. Scholz, and M. Mascher. 2017. MISA-web: a web server for microsatellite prediction. Bioinformatics 33: 2583–2585.

Bidartondo, M. I. 2005. The evolutionary ecology of myco-heterotrophy. New Phytologist 167: 335–352.

Bikaeff, J.-P. 2002. Liste des Orchidées de la région de Saül. Mouvement des Amis de la Nature pour le Groupement des Observations. Guyane française. 4 pp.

Bonen, L., and J. Vogel. 2001. The ins and outs of group II introns. Trends in Genetics 17: 322– 331.

Borba, E., S. MazzoniLViveiros, and J. Batista. 2014. Phylogenetic position and floral morphology of the Brazilian endemic, monospecific genus *Cotylolabium*: A sister group for the remaining Spiranthinae (Orchidaceae). Botanical Journal of the Linnean Society 175: 29–46.

Bromham, L., P. F. Cowman, and R. Lanfear. 2013. Parasitic plants have increased rates of molecular evolution across all three genomes. BMC Evolutionary Biology 13: 126.

Burnham, K. R., and D. R. Anderson. 2002. Model selection and multimodel inference: a practical information theoretic approach. Springer, New York.

Burns-Balogh, P., and H. Robinson. 1983. Evolution and phylogeny of the *Pelexia* alliance (Orchidaceae: Spiranthoideae: Spiranthinae). Systematic Botany 8: 263–268

Cai, L. 2023. Rethinking convergence in plant parasitism through the lens of molecular and population genetic processes. American Journal of Botany 110: e16174.

Chase, M. W., K. M. Cameron, J. V. Freudenstein, A. M. Pridgeon, G. Salazar, C. Van Den Berg, and A. Schuiteman. 2015. An updated classification of Orchidaceae. Botanical Journal of the Linnean Society 177: 151–174.

Chen, S., Y. Zhou, Y. Chen, and J. Gu. 2018. fastp: an ultra-fast all-in-one FASTQ preprocessor. Bioinformatics 34: i884–i890.

Cheuk, A., and T. Meier. 2021. Rotor subunits adaptations in ATP synthases from photosynthetic organisms. Biochemical Society Transactions 49: 541–550.

Crotty, S. M., B. Q. Minh, N. G. Bean, B. R. Holland, J. Tuke, L. S. Jermiin, and A. V. Haeseler. 2020. GHOST: Recovering historical signal from heterotachously evolved sequence alignments. Systematic Biology 69: 249–264.

Darling, A. E., B. Mau, and N. T. Perna. 2010. progressiveMauve: Multiple genome alignment with gene gain, loss and rearrangement. PLoS ONE 5: e11147.

Delannoy, E., S. Fujii, C., Colas Des Francs-Small, M. Brundrett, and I. Small. 2011. Rampant gene loss in the underground orchid *Rhizanthella gardneri* highlights evolutionary constraints on plastid genomes. Molecular Biology and Evolution 28: 2077–2086.

Determann, R. O. 1985. *Degranvillea* - a new orchid genus from French Guiana. American Orchid Society Bulletin. 54: 174.

Dierckxsens, N., P. Mardulyn, and G. Smits. 2017. NOVOPlasty: *de novo* assembly of organelle genomes from whole genome data. Nucleic Acids Research: gkw955.

Doyle, J. J. 2022. Defining coalescent genes: theory meets practice in organelle phylogenomics. Systematic Biology 71: 476–489.

Doyle, J.J. and Doyle, J.L. (1987) A rapid DNA isolation procedure for small quantities of fresh leaf tissue. Phytochemical Bulletin 19: 11–15

Drummond, A. J., and A. Rambaut. 2007. BEAST: Bayesian evolutionary analysis by sampling trees. BMC Evolutionary Biology 7: 214.

Dueck, L. A., D. Aygoren, and K. M. Cameron. 2014. A molecular framework for understanding the phylogeny of *Spiranthes* (Orchidaceae), a cosmopolitan genus with a North American center of diversity. American Journal of Botany 101: 1551–1571.

Dumont, V., E. Hágsater, and A. M. Pridgeon. 1996. Orchids: status survey and conservation action plan. IUCN. (last accessed 2023 Oct 30). https://portals.iucn.org/library/node/7026.

Freudenstein, J. V., and M. W. Chase. 2015. Phylogenetic relationships in Epidendroideae (Orchidaceae), one of the great flowering plant radiations: progressive specialization and diversification. Annals of Botany 115: 665–681.

Freudenstein, J. V., T. Yukawa, and Y. Luo. 2017. A reanalysis of relationships among Calypsoinae (Orchidaceae: Epidendroideae): floral and vegetative evolution and the placement of *Yoania*. Systematic Botany 42: 17–25.

Freudenstein, J.V. and Barrett, C.F. 2010. Mycoheterotrophy and diversity in Orchidaceae. Pp. 25–37 in: O. Seberg, G. Petersen, A. Barfod and J.I. Davis (eds.), Diversity, phylogeny and evolution in the monocotyledons. Aarhus: Aarhus University Press.

Funk, V., Hollowell, T., Berry, P., Kelloff, C. and S. N. Alexander. 2007. Checklist of the plants of the Guiana Shield (Venezuela: Amazonas, Bolivar, Delta Amacuro; Guyana, Surinam, French Guiana). Contributions from the United States National Herbarium 55: 1–580.

Garay, L. A. 1982. A generic revision of the Spiranthineae. Botanical Museum Leaflets, Harvard University. 28: 277–425

Garrett, N., J. Viruel, N. Klimpert, M. Soto Gomez, V. K. Y. Lam, V. S. F. T. Merckx, and S. W. Graham. 2023. Plastid phylogenomics and molecular evolution of Thismiaceae (Dioscoreales). American Journal of Botany 110: e16141.

GBIF.org. 2023. GBIF occurrence download 10.15468/dl.swcwyw (last accessed 11 October 2023).

Givnish, T. J., D. Spalink, M. Ames, S. P. Lyon, S. J. Hunter, A. Zuluaga, W. J. D. Iles, et al. 2015. Orchid phylogenomics and multiple drivers of their extraordinary diversification. *Proceedings*. Biological Sciences 282: 20151553.

Givnish, T. J., A. Zuluaga, D. Spalink, M. Soto Gomez, V. K. Y. Lam, J. M. Saarela, C. Sass, et al. 2018. Monocot plastid phylogenomics, timeline, net rates of species diversification, the power of multi-gene analyses, and a functional model for the origin of monocots. American Journal of Botany 105: 1888–1910.

Gorniak, M., J. Mytnik, P. Rutkowski, P. Tukałło, J. Minasiewicz, and D. Szlachetko. 2006. Phylogenetic relationships within the subtribe Spiranthinae s.l. (Orchidaceae) inferred from the nuclear ITS region. *Biodiversity*, Research and Conservation 1–2: 18–24.

Graham, S. W., V. K. Y. Lam, and V. S. F. T. Merckx. 2017. Plastomes on the edge: the evolutionary breakdown of mycoheterotroph plastid genomes. The New Phytologist 214: 48–55.

Guo, X., Z. Zhao, S. S. Mar, D. Zhang, and R. M. K. Saunders. 2019. A symbiotic balancing act: arbuscular mycorrhizal specificity and specialist fungus gnat pollination in the mycoheterotrophic genus *Thismia* (Thismiaceae). Annals of Botany 124: 331–342.

Hertweck, K. L., M. S. Kinney, S. A. Stuart, O. Maurin, S. Mathews, M. W. Chase, M. A. Gandolfo, and J. C. Pires. 2015. Phylogenetics, divergence times and diversification from three genomic partitions in monocots. Botanical Journal of the Linnean Society 178: 375– 393.

Jansen, R. K., and T. A. Ruhlman. 2012. Plastid Genomes of Seed Plants. In R. Bock, and V. Knoop [eds.], Genomics of Chloroplasts and Mitochondria, Advances in Photosynthesis and Respiration, 103–126. Springer Netherlands, Dordrecht.

Jin, J.-J., W.-B. Yu, J.-B. Yang, Y. Song, C. W. dePamphilis, T.-S. Yi, and D.-Z. Li. 2020. GetOrganelle: a fast and versatile toolkit for accurate de novo assembly of organelle genomes. Genome Biology 21: 241.

Jombart, T., and S. Dray. 2008. adephylo: exploratory analyses for the phylogenetic comparative method. Bioinformatics 26: 1907–1909.

Jones, B. R., and A. F. Y. Poon. 2017. node.dating: dating ancestors in phylogenetic trees in R. Bioinformatics 33: 932–934.

Jónsson, H., A. Ginolhac, M. Schubert, P. L. F. Johnson, and L. Orlando. 2013. mapDamage2.0: fast approximate Bayesian estimates of ancient DNA damage parameters. Bioinformatics 29: 1682–1684.

Kalyaanamoorthy, S., B. Q. Minh, T. K. F. Wong, A. von Haeseler, and L. S. Jermiin. 2017. ModelFinder: fast model selection for accurate phylogenetic estimates. Nature Methods 14: 587–589.

Kamikawa, R., G. Tanifuji, S. A. Ishikawa, K.-I. Ishii, Y. Matsuno, N. T. Onodera, K.-I. Ishida, et al. 2015. Proposal of a twin arginine translocator system-mediated constraint against loss of ATP synthase genes from nonphotosynthetic plastid genomes. Molecular Biology and Evolution 32: 2598–2604.

Katoh, K., and D. M. Standley. 2013. MAFFT multiple sequence alignment software version 7: improvements in performance and usability. Molecular Biology and Evolution 30: 772–780.

Kent, W. J. 2002. BLAT—the BLAST-like alignment tool. Genome Research 12: 656–664.

Kim, Y.-K., S. Jo, S.-H. Cheon, M.-J. Joo, J.-R. Hong, M. H. Kwak, and K.-J. Kim. 2019. Extensive losses of photosynthesis genes in the plastome of a mycoheterotrophic orchid, *Cyrtosia septentrionalis* (Vanilloideae: Orchidaceae). Genome Biology and Evolution 11: 565–571.

Kim, Y.-K., S. Jo, S.-H. Cheon, M.-J. Joo, J.-R. Hong, M. Kwak, and K.-J. Kim. 2020. Plastome evolution and phylogeny of Orchidaceae, with 24 new sequences. Frontiers in Plant Science 11.

Kim, W., T. Lautenschläger, J. F. Bolin, M. Rees, A. Nzuzi, R. Zhou, S. Wanke, and M. Jost. 2023. Extreme plastomes in holoparasitic Balanophoraceae are not the norm. BMC Genomics 24: 330.

Klimpert, N. J., J. L. S. Mayer, D. S. Sarzi, F. Prosdocimi, F. Pinheiro, and S. W. Graham. 2022. Phylogenomics and plastome evolution of a Brazilian mycoheterotrophic orchid, *Pogoniopsis schenckii*. American Journal of Botany 109: 2030–2050.

Knox, E. B. 2014. The dynamic history of plastid genomes in the Campanulaceae *sensu lato* is unique among angiosperms. Proceedings of the National Academy of Sciences 111: 11097– 11102.

Knox, E. B. 2022. DNA sequence analysis of an inversion hot spot in Lobeliaceae plastomes. Plants 11: 2863.

Kolde, R. 2019. pheatmap: Pretty heatmaps. R package version 1.0.12. https://CRAN.R-project.org/package=pheatmap.

Kosakovsky Pond, S. L., S. D. W. Frost, and S. V. Muse. 2005. HyPhy: hypothesis testing using phylogenies. Bioinformatics 21: 676–679.

Kumar, S., and S. B. Hedges. 2016. Advances in time estimation methods for molecular data. Molecular Biology and Evolution 33: 863–869.

Lam, V. K. Y., M. Soto Gomez, and S. W. Graham. 2015. The highly reduced plastome of mycoheterotrophic *Sciaphila* (Triuridaceae) is colinear with its green relatives and is under strong purifying selection. Genome Biology and Evolution 7: 2220–2236.

Lam, V. K. Y., H. Darby, V. S. F. T. Merckx, G. Lim, T. Yukawa, K. M. Neubig, J. R. Abbott, et al. 2018. Phylogenomic inference *in extremis*: A case study with mycoheterotroph plastomes. American Journal of Botany 105: 480–494.

Lartillot, N., and R. Poujol. 2011. A phylogenetic model for investigating correlated evolution of substitution rates and continuous phenotypic characters. Molecular Biology and Evolution 28: 729–744.

Leake, J. R. 1994. The biology of myco-heterotrophic (‘saprophytic’) plants. The New Phytologist 127: 171–216.

Liere, K., A. Weihe, and T. Börner. 2011. The transcription machineries of plant mitochondria and chloroplasts: Composition, function, and regulation. Journal of Plant Physiology 168: 1345–1360.

Menéndez, C. D., P. Poczai, B. Williams, L. Myllys, and A. Amiryousefi. 2023. IRplus: An augmented tool to detect inverted repeats in plastid genomes. Genome Biology and Evolution 15.

Merckx, V. S. F. T., and E. F. Smets. 2014. *Thismia americana*, the 101st anniversary of a botanical mystery. International Journal of Plant Sciences 175: 165–175.

Merckx, V. S. F. T., J. V. Freudenstein, J. Kissling, M. J. M. Christenhusz, R. E. Stotler, B. Crandall-Stotler, N. Wickett, et al. 2013a. Taxonomy and Classification. In V. Merckx [ed.], Mycoheterotrophy: The Biology of Plants Living on Fungi, 19–101. Springer, New York, NY.

Merckx, V.S.F.T., E.F. Smets, and C.D. Specht. 2013b. Biogeography and Conservation. In: Merckx V.S.F.T. (ed.). Mycoheterotrophy: The Biology of Plants Living on Fungi. Springer-Verlag, New York, New York.

Merckx, V. S. F. T., S. I. F. Gomes, M. Wapstra, C. Hunt, G. Steenbeeke, C. B. Mennes, N. Walsh, et al. 2017. The biogeographical history of the interaction between mycoheterotrophic *Thismia* (Thismiaceae) plants and mycorrhizal *Rhizophagus* (Glomeraceae) fungi. Journal of Biogeography 44: 1869–1879.

Microsoft Corporation. 2018. Microsoft Excel. Retrieved from https://office.microsoft.com/excel.

Minh, B. Q., H. A. Schmidt, O. Chernomor, D. Schrempf, M. D. Woodhams, A. Von Haeseler, and R. Lanfear. 2020. IQ-TREE 2: New models and efficient methods for phylogenetic inference in the genomic era. Molecular Biology and Evolution 37: 1530–1534.

Miura, S., K. Tamura, Q. Tao, L. A. Huuki, S. L. Kosakovsky Pond, J. Priest, J. Deng, and S. Kumar. 2020. A new method for inferring timetrees from temporally sampled molecular sequences. PLOS Computational Biology 16: e1007046.

Molina, J., K. M. Hazzouri, D. Nickrent, M. Geisler, R. S. Meyer, M. M. Pentony, J. M. Flowers, et al. 2014. Possible loss of the chloroplast genome in the parasitic flowering plant *Rafflesia lagascae* (Rafflesiaceae). Molecular Biology and Evolution 31: 793–803.

Motomura, H., M.-A. Selosse, F. Martos, A. Kagawa, and T. Yukawa. 2010. Mycoheterotrophy evolved from mixotrophic ancestors: evidence in *Cymbidium* (Orchidaceae). Annals of Botany 106: 573–581.

Mower, J. P., and T. L. Vickrey. 2018. Chapter Nine - Structural diversity among plastid genomes of land plants. In S.-M. Chaw, and R. K. Jansen [eds.], Advances in Botanical Research, Plastid Genome Evolution, 263–292. Academic Press.

Murray, K. J. H. 2019. Chloroplast genome evolution in New Zealand mycoheterotrophic Orchidaceae: a thesis presented in partial fulfilment of the requirements for the degree of Master of Science in Plant Biology at Massey University, Manawatu, New Zealand. Massey University.

Naumann, J., J. P. Der, E. K. Wafula, S. S. Jones, S. T. Wagner, L. A. Honaas, P. E. Ralph, et al. 2016. Detecting and characterizing the highly divergent plastid genome of the nonphotosynthetic parasitic plant *Hydnora visseri* (Hydnoraceae). Genome Biology and Evolution 8: 345–363.

New, C. P., Q. Ma, and C. Dabney-Smith. 2018. Routing of thylakoid lumen proteins by the chloroplast twin arginine transport pathway. Photosynthesis Research 138: 289–301.

O’Byrne, P. 2014. On the evolution of *Dipodium* R. BR. Reinwardtia. 14: 123–132.

Ogura-Tsujita, Y., T. Yukawa, and A. Kinoshita. 2021. Evolutionary histories and mycorrhizal associations of mycoheterotrophic plants dependent on saprotrophic fungi. Journal of Plant Research 134: 19–41.

Paradis, E., and K. Schliep. 2019. ape 5.0: an environment for modern phylogenetics and evolutionary analyses in R. Bioinformatics 35: 526–528.

Pelser, P. B., D. L. Nickrent, B. W. Van Ee, and J. F. Barcelona. 2019. A phylogenetic and biogeographic study of *Rafflesia* (Rafflesiaceae) in the Philippines: Limited dispersal and high island endemism. Molecular Phylogenetics and Evolution 139: 106555.

Peltier, G., E.-M. Aro, and T. Shikanai. 2016. NDH-1 and NDH-2 Plastoquinone Reductases in oxygenic photosynthesis. Annual Review of Plant Biology 67: 55–80.

Pennell, M., Eastman, J., Slater, G., Brown, J., Uyeda, J., Fitzjohn R, Alfaro, M., et al. 2014. geiger v2.0: an expanded suite of methods for fitting macroevolutionary models to phylogenetic trees. Bioinformatics 30: 2216–2218.

Perez-Escobar, O. A., D. Bogarín, N. A. S. Przelomska, J. D. Ackerman, J. A. Balbuena, S. Bellot, R. P. Bühlmann, et al. 2023. The origin and speciation of orchids. bioRxiv 10.1101/2023.09.10.556973.

Perry, A. S., and K. H. Wolfe. 2002. Nucleotide substitution rates in legume chloroplast DNA depend on the presence of the inverted repeat. Journal of Molecular Evolution 55: 501–508.

Pridgeon, A. M., P. J. Cribb, M. W. Chase, and F. N. Rasmussen [eds.]. 2001. Genera Orchidacearum: Volume 2: Orchidoideae (Part 1). Oxford University Press, Oxford, New York.

Pridgeon, E. by A. M., P. J. Cribb, M. W. Chase, and F. N. Rasmussen [eds.]. 2003. Genera Orchidacearum: Volume 3: Orchidoideae (Part 2), Vanilloideae. Oxford University Press, Oxford, New York.

R Core Team. 2023. R: A language and environment for statistical computing. R Foundation for Statistical Computing, Vienna, Austria. https://www.R-project.org/.

Rambaut, A. 2010. FigTree v1.3.1. Institute of Evolutionary Biology, University of Edinburgh, Edinburgh. http://tree.bio.ed.ac.uk/software/figtree.

Ramírez, S. R., B. Gravendeel, R. B. Singer, C. R. Marshall, and N. E. Pierce. 2007. Dating the origin of the Orchidaceae from a fossil orchid with its pollinator. Nature 448: 1042–1045.

Ranwez, V., E. J. P. Douzery, C. Cambon, N. Chantret, and F. Delsuc. 2018. MACSE v2: Toolkit for the alignment of coding sequences accounting for frameshifts and stop codons. Molecular Biology and Evolution 35: 2582–2584.

Rasmussen, H. N. 1995. Terrestrial orchids: from seed to mycotrophic plant. Cambridge University Press.

Rasmussen, H. N., K. W. Dixon, J. Jersáková, and T. Těšitelová. 2015. Germination and seedling establishment in orchids: a complex of requirements. Annals of Botany 116: 391–402.

Rice P., Longden, I., and A. Bleasby. 2000. EMBOSS: The European Molecular Biology Open Software Suite. Trends in Genetics 16: 276–277.

Rice, P., I. Longden, and A. Bleasby. 2000. EMBOSS: The European Molecular Biology Open Software Suite. Trends in Genetics 16: 276–277.

Ruhlman, T. A., and R. K. Jansen. 2014. The plastid genomes of flowering plants. Methods in Molecular Biology 1132: 3–38.

Salazar, G. A., M. W. Chase, M. A. Soto Arenas, and M. Ingrouille. 2003. Phylogenetics of Cranichideae with emphasis on Spiranthinae (Orchidaceae, Orchidoideae): evidence from plastid and nuclear DNA sequences. American Journal of Botany 90: 777–795.

Salazar, G. A., J. A. N. Batista, L. I. Cabrera, C. van Den Berg, W. M. Whitten, E. C. Smidt, C. R. Buzatto, et al. 2018. Phylogenetic systematics of subtribe Spiranthinae (Orchidaceae: Orchidoideae: Cranichideae) based on nuclear and plastid DNA sequences of a nearly complete generic sample. Botanical Journal of the Linnean Society 186: 273–303.

Salazar, G. A., L. Baquero, M. Jiménez, and F. Rizo-Patrón. 2022. DNA links Andean tepui endemic *Helonoma peruvianato* to *Hapalorchis* (Orchidaceae, Spiranthinae. Phytotaxa 574: 61–72.

Sanchez-Puerta, M. V., L. F. Ceriotti, L. M. Gatica-Soria, M. E. Roulet, L. E. Garcia, and H. A. Sato. 2023. Beyond parasitic convergence: unravelling the evolution of the organellar genomes in holoparasites. Annals of Botany: mcad108.

Schelkunov, M. I., V. Y. Shtratnikova, M. S. Nuraliev, M.-A. Selosse, A. A. Penin, and M. D. Logacheva. 2015. Exploring the limits for reduction of plastid genomes: a case study of the mycoheterotrophic orchids *Epipogium aphyllum* and *Epipogium roseum*. Genome Biology and Evolution 7: 1179–1191.

Schwarz, G. E. 1978. Estimating the dimension of a model. Annals of Statistics 6: 461–464.

Serna-Sánchez, M. A., O. A. Pérez-Escobar, D. Bogarín, M. F. Torres-Jimenez, A. C. Alvarez-Yela, J. E. Arcila-Galvis, C. F. Hall, et al. 2021. Plastid phylogenomics resolves ambiguous relationships within the orchid family and provides a solid timeframe for biogeography and macroevolution. Scientific Reports 11: 6858.

Shepeleva, E., M. Schelkunov, M. Hroneš, M. Sochor, M. Dančák, M. V.S.F.T., K. I.A.B.S., et al. 2020. Phylogenetics of the mycoheterotrophic genus *Thismia* (Thismiaceae: Dioscoreales) with a focus on the Old-World taxa: delineation of novel natural groups and insights into the evolution of morphological traits. Botanical Journal of the Linnean Society 193: 287–315.

Sinn, B. T., D. D. Sedmak, L. M. Kelly, and J. V. Freudenstein. 2018. Total duplication of the small single copy region in the angiosperm plastome: Rearrangement and inverted repeat instability in *Asarum*. American Journal of Botany 105: 71–84.

Smith, D. R., and R. W. Lee. 2014. A plastid without a genome: evidence from the nonphotosynthetic green algal genus *Polytomella*. Plant Physiology 164: 1812–1819.

Steyermark, J. A., and B. K. Holst. 1989. Flora of the Venezuelan Guayana–VII contributions to the flora of the Cerro Aracamuni, Venezuela. Annals of the Missouri Botanical Garden 76: 945–992.

Taylor, D. L., C. F. Barrett, G. E. Beatty, S. E. Hopkins, A. H. Kennedy, and M. R. Klooster. 2013. Progress and prospects for the ecological genetics of mycoheterotrophs. In V. Merckx [ed.], Mycoheterotrophy: The Biology of Plants Living on Fungi, 245–266. Springer, New York, NY.

Těšitel, J. 2016. Functional biology of parasitic plants: a review. Plant Ecology and Evolution 149: 5–20.

Tillich, M., P. Lehwark, T. Pellizzer, E. S. Ulbricht-Jones, A. Fischer, R. Bock, and S. Greiner. 2017. GeSeq – versatile and accurate annotation of organelle genomes. Nucleic Acids Research 45: W6–W11.

Timilsena, P. R., C. F. Barrett, A. Piñeyro-Nelson, E. K. Wafula, S. Ayyampalayam, J. R. McNeal, T. Yukawa, et al. 2023. Phylotranscriptomic Analyses of mycoheterotrophic monocots show a continuum of convergent evolutionary changes in expressed nuclear genes from three independent nonphotosynthetic lineages. Genome Biology and Evolution 15: evac183.

To, T.-H., M. Jung, S. Lycett, and O. Gascuel. 2016. Fast dating using least-squares criteria and algorithms. Systematic Biology 65: 82–97.

Tu, X.-D., D.-K. Liu, S.-W. Xu, C.-Y. Zhou, X.-Y. Gao, M.-Y. Zeng, S. Zhang, et al. 2021. Plastid phylogenomics improves resolution of phylogenetic relationship in the *Cheirostylis* and *Goodyera* clades of Goodyerinae (Orchidoideae, Orchidaceae). Molecular Phylogenetics and Evolution 164: 107269.

Twyford, A. D. 2018. Parasitic plants. Current Biology 28: R857–R859.

Twyford, A. D., and R. W. Ness. 2017. Strategies for complete plastid genome sequencing. Molecular Ecology Resources 17: 858–868.

Véron, S., Rodrigues-Vaz, C., Lebreton, E., Ah-Peng, C., Boullet, V., Chevillotte, H., Gradstein, S., et al. 2021. An assessment of the endemic spermatophytes, pteridophytes and bryophytes of the French Overseas Territories: towards a better conservation outlook. Biodiversity and Conservation 30: 2097–2124.

Vogel, J., T. Börner, and W. R. Hess. 1999. Comparative analysis of splicing of the complete set of chloroplast group II introns in three higher plant mutants. Nucleic Acids Research 27: 3866–3874.

Wang, Z.-X., D.-J. Wang, and T.-S. Yi. 2022. Does IR-loss promote plastome structural variation and sequence evolution? Frontiers in Plant Science 13: 888049.

Waterman, R. J., and M. I. Bidartondo. 2008. Deception above, deception below: linking pollination and mycorrhizal biology of orchids. Journal of Experimental Botany 59: 1085– 1096.

Wen, Y., Y. Qin, B. Shao, J. Li, C. Ma, Y. Liu, B. Yang, and X. Jin. 2022. The extremely reduced, diverged and reconfigured plastomes of the largest mycoheterotrophic orchid lineage. BMC Plant Biology 22: 448.

Wertheim, J. O., B. Murrell, M. D. Smith, S. L. Kosakovsky Pond, and K. Scheffler. 2015. RELAX: Detecting relaxed selection in a phylogenetic framework. Molecular Biology and Evolution 32: 820–832.

Westwood, J. H., J. I. Yoder, M. P. Timko, and C. W. dePamphilis. 2010. The evolution of parasitism in plants. Trends in Plant Science 15: 227–235.

Wicke, S., and J. Naumann. 2018. Chapter Eleven - Molecular Evolution of Plastid Genomes in Parasitic Flowering Plants. In S.-M. Chaw, and R. K. Jansen [eds.], Advances in Botanical Research, Plastid Genome Evolution, 315–347. Academic Press.

Wicke, S., G. M. Schneeweiss, C. W. dePamphilis, K. F. Müller, and D. Quandt. 2011. The evolution of the plastid chromosome in land plants: gene content, gene order, gene function. Plant Molecular Biology 76: 273–297.

Wicke, S., K. F. Müller, C. W. dePamphilis, D. Quandt, S. Bellot, and G. M. Schneeweiss. 2016. Mechanistic model of evolutionary rate variation en route to a nonphotosynthetic lifestyle in plants. Proceedings of the National Academy of Sciences 113: 9045–9050.

Wickett, N. J., S. Mirarab, N. Nguyen, T. Warnow, E. Carpenter, N. Matasci, S. Ayyampalayam, et al. 2014. Phylotranscriptomic analysis of the origin and early diversification of land plants. Proceedings of the National Academy of Sciences 111: E4859–E4868.

Wolfe, K. H., W. H. Li, and P. M. Sharp. 1987. Rates of nucleotide substitution vary greatly among plant mitochondrial, chloroplast, and nuclear DNAs. Proceedings of the National Academy of Sciences 84: 9054–9058.

Yuan, Y., X. Jin, J. Liu, X. Zhao, J. Zhou, X. Wang, D. Wang, et al. 2018. The *Gastrodia elata* genome provides insights into plant adaptation to heterotrophy. Nature Communications 9: 1615.

Yudina, S. V., M. I. Schelkunov, L. Nauheimer, D. Crayn, S. Chantanaorrapint, M. Hroneš, M. Sochor, et al. 2021. Comparative analysis of plastid genomes in the non-photosynthetic genus *Thismia* reveals ongoing gene set reduction. Frontiers in Plant Science 12: 602598.

Zhang, D., F. Gao, I. Jakovlić, H. Zou, J. Zhang, W. X. Li, and G. T. Wang. 2020. PhyloSuite: An integrated and scalable desktop platform for streamlined molecular sequence data management and evolutionary phylogenetics studies. Molecular Ecology Resources 20: 348–355.

Zhang, G., Y. Hu, M.-Z. Huang, W.-C. Huang, D.-K. Liu, D. Zhang, H. Hu, et al. 2023. Comprehensive phylogenetic analyses of Orchidaceae using nuclear genes and evolutionary insights into epiphytism. Journal of Integrative Plant Biology 65: 1204–1225.

Zhong, X. 2020. Assembly, Annotation and Analysis of Chloroplast Genomes. Available online: https://research-repository.uwa.edu.au/en/publications/assembly-annotation-and-analysis-of-chloroplast-genomes (accessed on 31 October 2023).

Zoschke, R., M. Nakamura, K. Liere, M. Sugiura, T. Börner, and C. Schmitz-Linneweber. 2010. An organellar maturase associates with multiple group II introns. Proceedings of the National Academy of Sciences 107: 3245–3250.

Zuker, M. 2003. Mfold web server for nucleic acid folding and hybridization prediction. Nucleic Acids Research 31: 3406–3415.

