## Supplementary Information for "Plastid genome evolution in leafless members of the orchid subfamily Orchidoideae, with a focus on *Degranvillea dermaptera*"

### Barrett et al.—American Journal of Botany 2024 – Appendix S1
### Postmortem DNA substitutions for *Degranvillea*

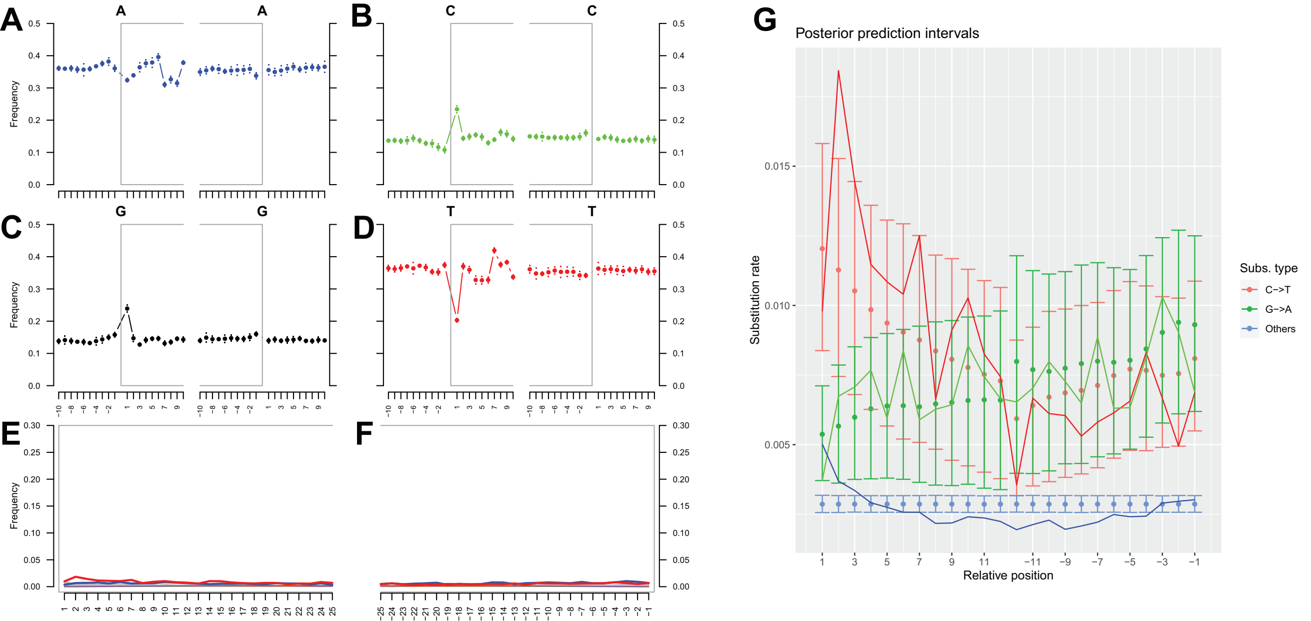

Barrett et al.—American Journal of Botany 2024 – Appendix S2
Inverted repeat dynamics in leafless orchidoid orchids

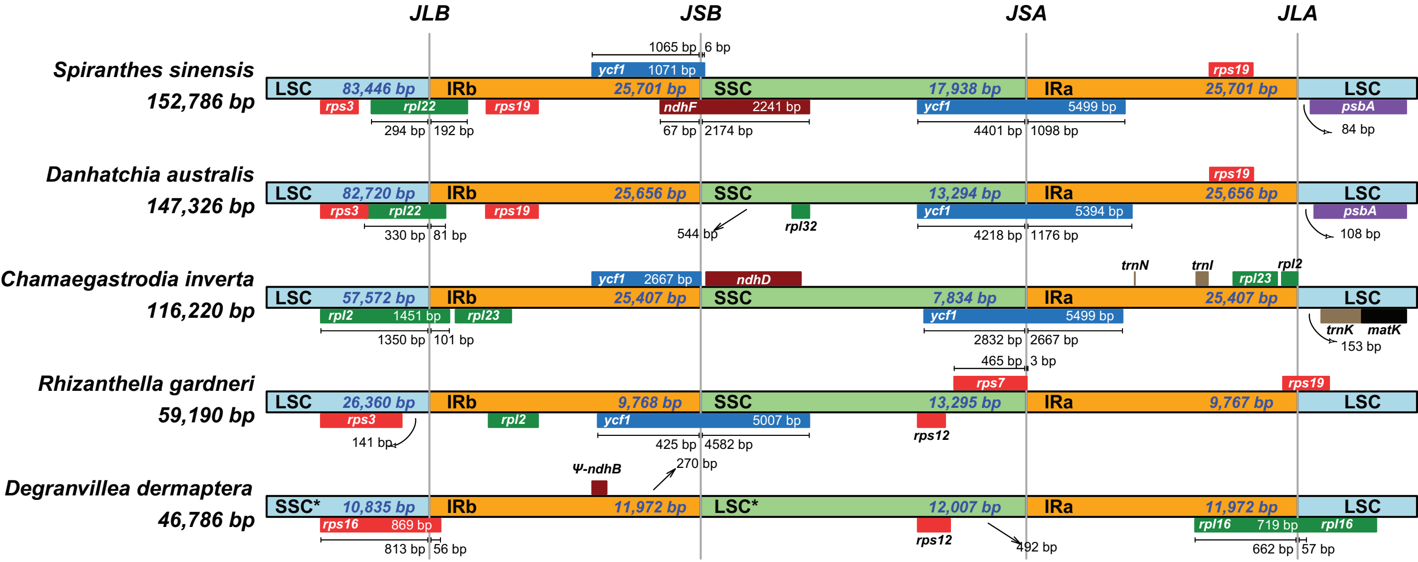

Barrett et al.—American Journal of Botany 2024 – Appendix S3

### Palindromic repeats near inversion breakpoints of *Degranvillea*

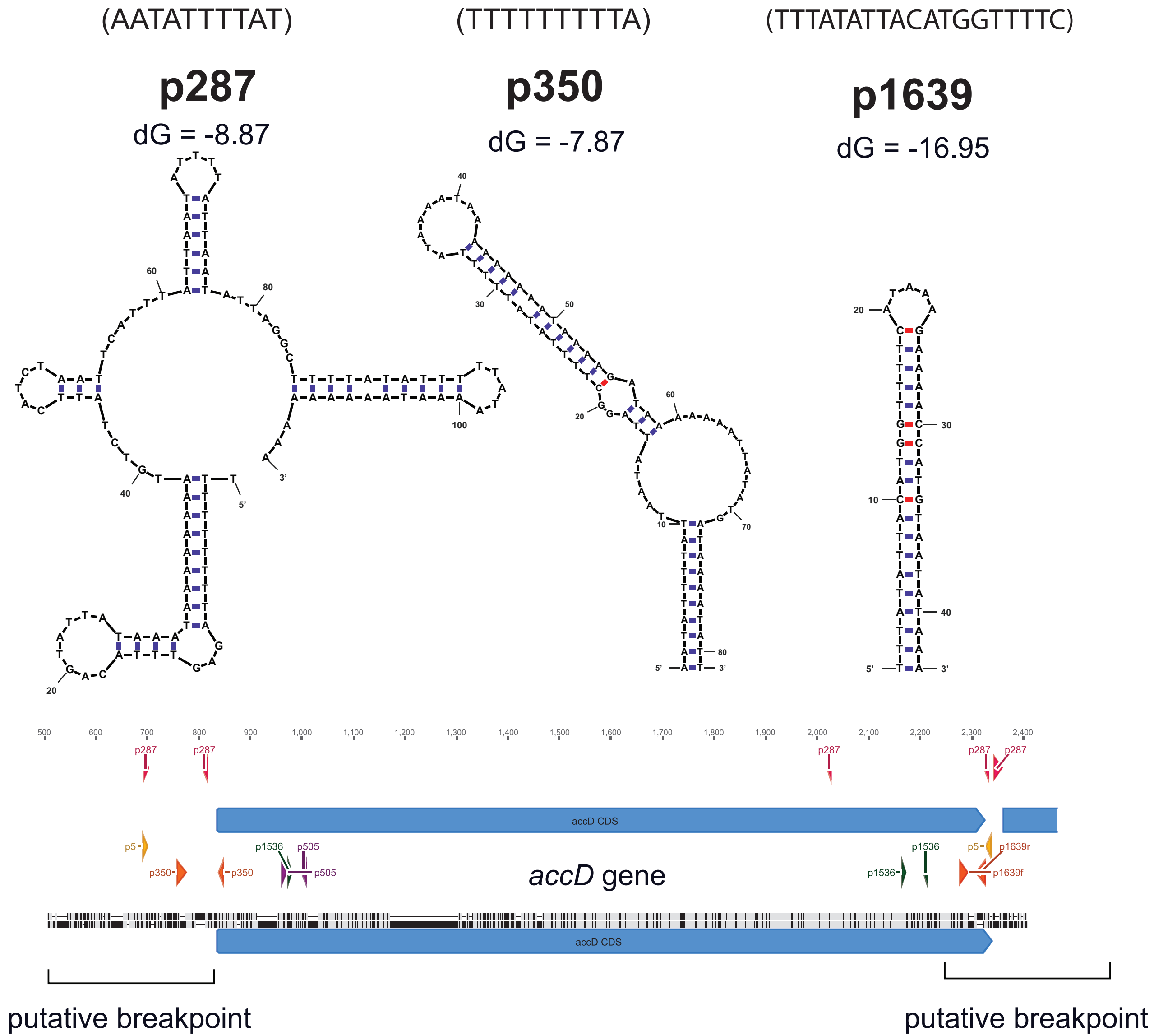

Barrett et al.—American Journal of Botany 2024 – Appendix S4
Plastid least-squares divergence time estimates for Orchidoideae

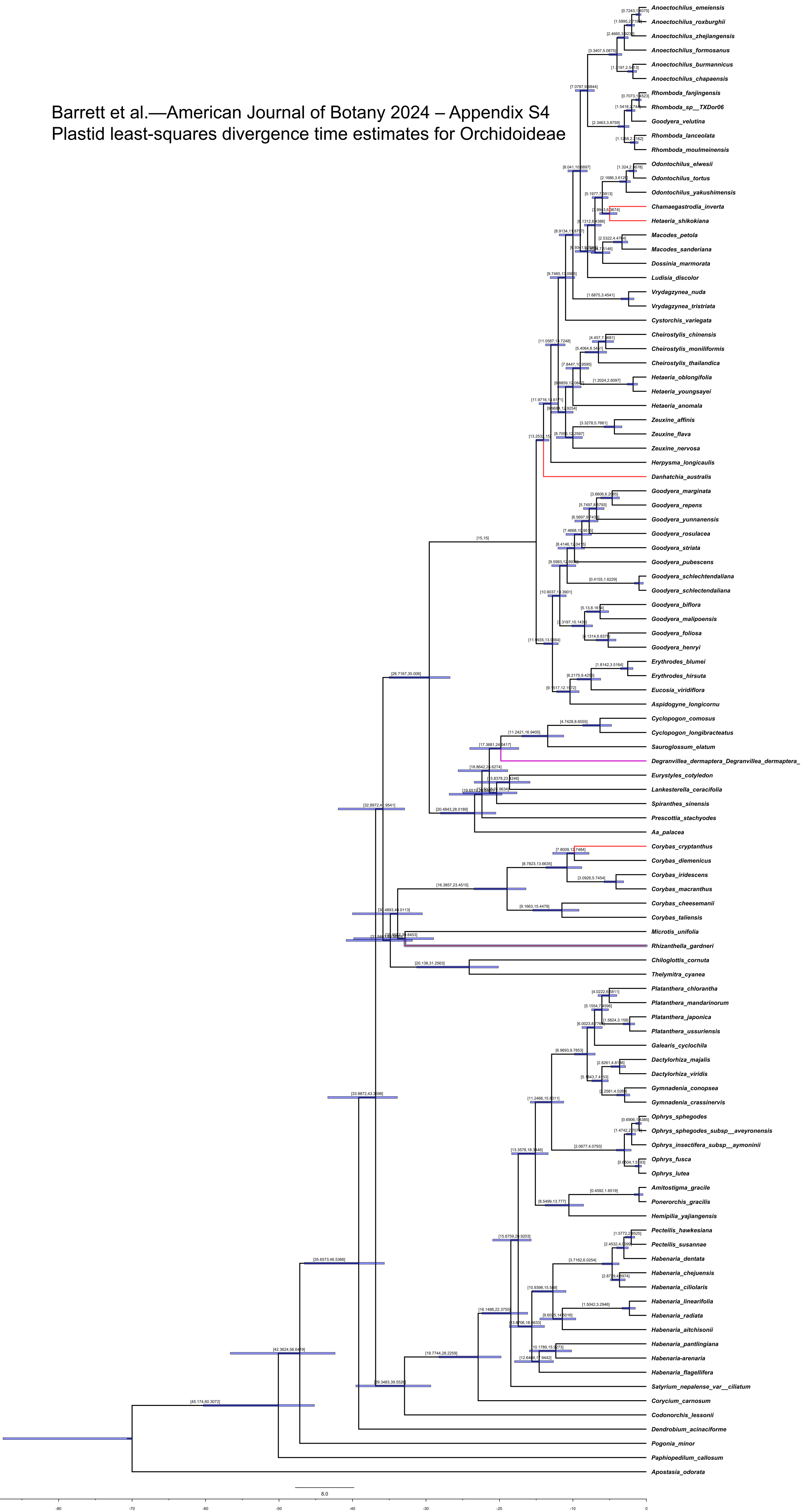

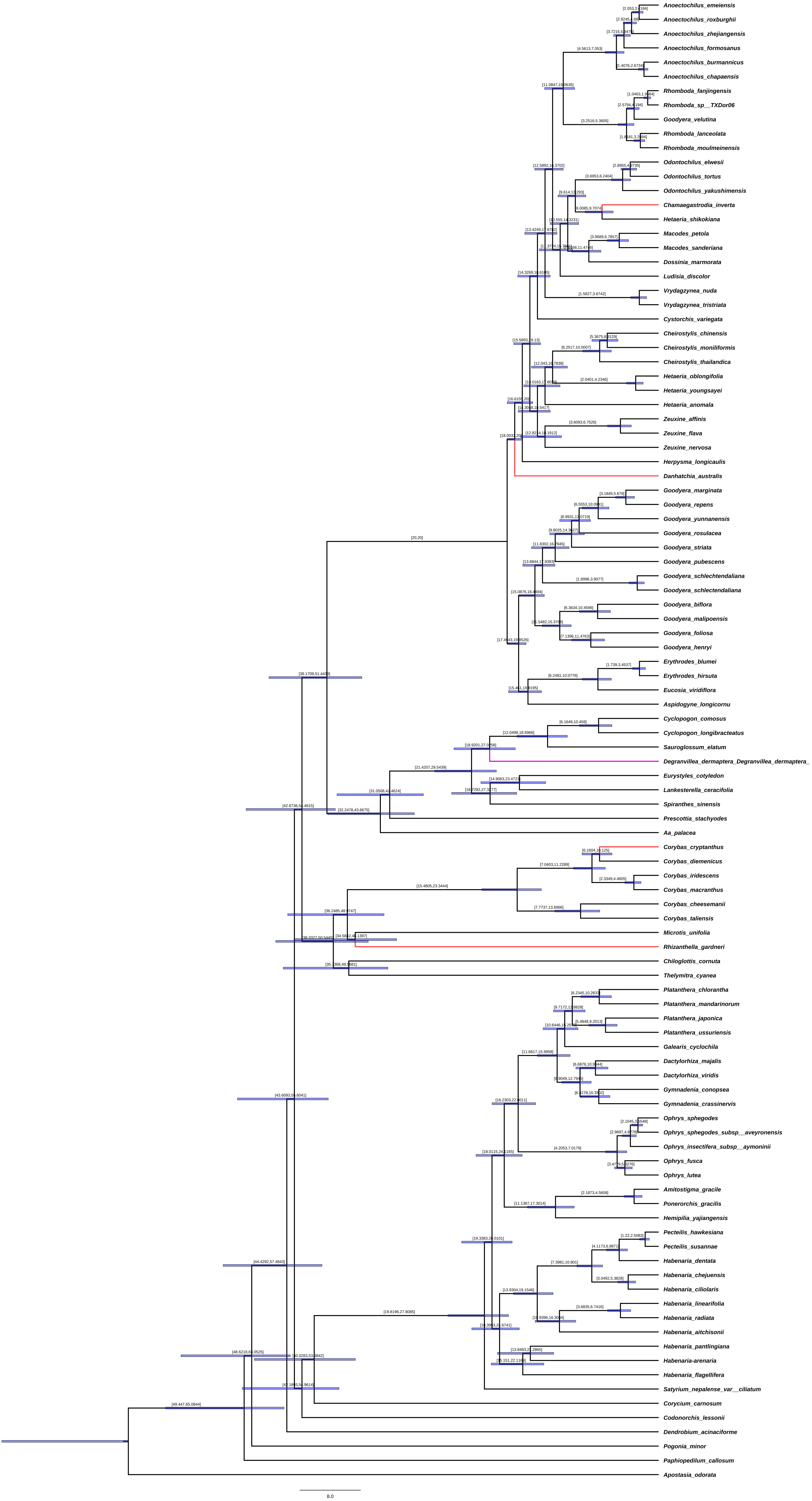

Anoectochilus_emeiensis
Anoectochilus_roxburghii
Anoectochilus_zhejiangensis
Anoectochilus_formosanus
Anoectochilus_burmannicus
Anoectochilus_chapaensis
Rhomboda_fanjingensis
Rhomboda_sp__TXDor06
Goodyera_velutina
Rhomboda_lanceolata
Rhomboda_moulmeinensis
Odontochilus_elwesii
Odontochilus_tortus
Odontochilus_yakushimensis
Chamaegastrodia_inverta
Hetaeria_shikokiana
Macodes_petola
Macodes_sanderiana
Dossinia_marmorata
Ludisia_discolor
Vrydagzynea_nuda
Vrydagzynea_tristriata
Cystorchis_variegata
Cheirostylis_chinensis
Cheirostylis_moniliformis
Cheirostylis_thailandica
Hetaeria_oblongifolia
Hetaeria_youngsayei
Hetaeria_anomala
Zeuxine_affinis
Zeuxine_flava
Zeuxine_nervosa
Herpysma_longicaulis
Danhatchia_australis
Goodyera_marginata
Goodyera_repens
Goodyera_yunnanensis
Goodyera_rosulacea
Goodyera_striata
Goodyera_pubescens
Goodyera_schlechtendaliana
Goodyera_schlectendaliana
Goodyera_biflora
Goodyera_malipoensis
Goodyera_foliosa
Goodyera_henryi
Erythrodes_blumei
Erythrodes_hirsuta
Eucosia_viridiflora
Aspidogyne_longicornu
Cyclopogon_comosus
Cyclopogon_longibracteatus
Sauroglossum_elatum
Degranvillea_dermaptera_Degranvillea_dermaptera_
Eurystyles_cotyledon
Lankesterella_ceracifolia
Spiranthes_sinensis
Prescottia_stachyodes
Aa_palacea
Corybas_cryptanthus
Corybas_diemenicus
Corybas_iridescens
Corybas_macranthus
Corybas_cheesemanii
Corybas_taliensis
Microtis_unifolia
Rhizanthella_gardneri
Chiloglottis_cornuta
Thelymitra_cyanea
Platanthera_chlorantha
Platanthera_mandarinorum
Platanthera_japonica
Platanthera_ussuriensis
Galearis_cyclochila
Dactylorhiza_majalis
Dactylorhiza_viridis
Gymnadenia_conopsea
Gymnadenia_crassinervis
Ophrys_sphegodes
Ophrys_sphegodes_subsp__aveyronensis
Ophrys_insectifera_subsp__aymoninii
Ophrys_fusca
Ophrys_lutea
Amitostigma_gracile
Ponerorchis_gracilis
Hemipilia_yajiangensis
Pecteilis_hawkesiana
Pecteilis_susannae
Habenaria_dentata
Habenaria_chejuensis
Habenaria_ciliolaris
Habenaria_linearifolia
Habenaria_radiata
Habenaria_aitchisonii
Habenaria_pantlingiana
Habenaria-arenaria
Habenaria_flagellifera
Satyrium_nepalense_var__ciliatum
Corycium_carnosum
Codonorchis_lessonii
Dendrobium_acinaciforme
Pogonia_minor
Paphiopedilum_callosum
Apostasia_odorata
8.0

Barrett et al.—American Journal of Botany 2024 – Appendix S5
ITS least-squares divergence times for Spiranthinae

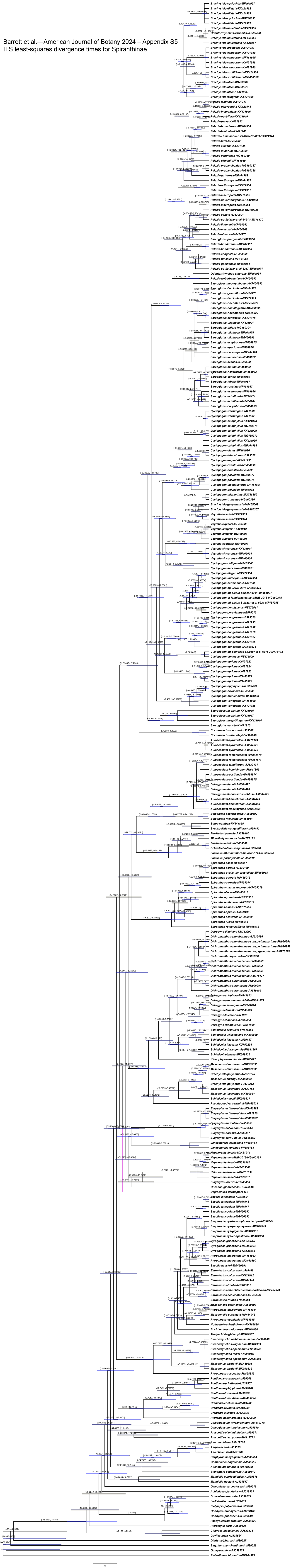

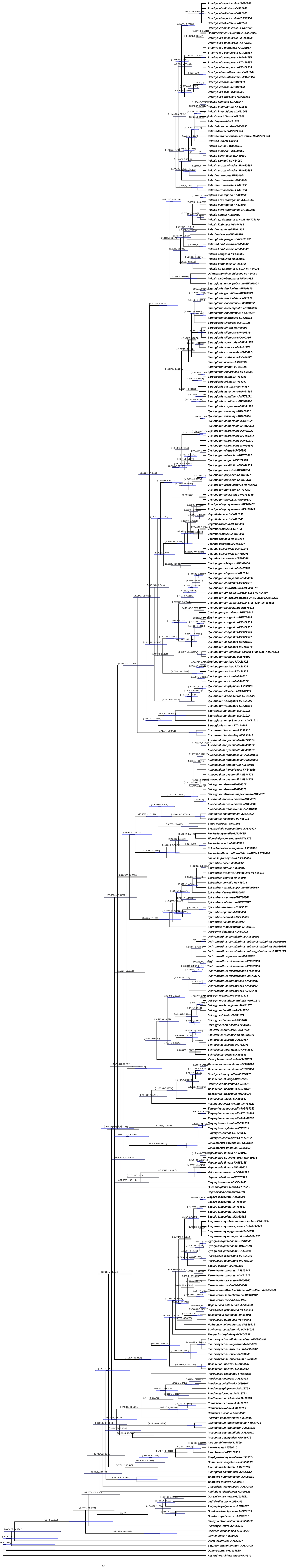

Barrett et al.—*American Journal of Botany* 2024 – Appendix S6

Intron lengths (bp) among leafless orchidoid orchids and the leafy *Spiranthes sinensis*.

| | ***Spiranthes sinensis*** | ***Danhatchia australis*** | ***Chamae. inverta*** | ***Corybas cryptanthus*** | ***Rhizanthella gardneri*** | ***Degranvillea dermaptera*** |
|---|---|---|---|---|---|---|
| ***atpF**** | 772 | 924 | | | | |
| ***clpP intron 2**** | 608 | 640 | 653 | 526 | 635 | 641 |
| ***clpP intron 1*** | 929 | 928 | 843 | 783 | 817 | 674 |
| ***ndhB*** | 704 | 130 | | | | |
| ***petB*** | 744 | 738 | | | | |
| ***petD*** | 819 | 778 | | | | |
| ***rpl2**** | 669 | 664 | 671 | 579 | 582 | 669 |
| ***rpl16*** | 1197 | 1143 | 1267 | 900 | 1027 | 820 |
| ***rpoC1*** | 741 | 734 | | | | |
| ***rps16*** | | 909 | 756 | 673 | | 624 |
| ***rps12-3'**** | 534 | 542 | 550 | 552 | | 636 |
| ***trnA-UGC**** | 798 | 804 | 815 | | | |
| ***trnG-UCC*** | 670 | 673 | 679 | 565 | | |
| ***trnI-GAU**** | 935 | 936 | 953 | | | |
| ***trnL-UAA*** | 646 | 602 | 593 | 353 | | |
| ***trnV-UAC*** | 600 | 587 | 583 | | | |
| ***trnK-UUU*-matK*** | 278 | 231 | 233 | 135 | | |
| ***matK-trnK-UUU**** | 1081 | 1091 | 1083 | 789 | | |
